# Nonlinear spatial integration underlies the diversity of retinal ganglion cell responses to natural stimuli

**DOI:** 10.1101/2020.06.18.159384

**Authors:** Dimokratis Karamanlis, Tim Gollisch

## Abstract

How neurons encode natural stimuli is a fundamental question for sensory neuroscience. In the early visual system, standard encoding models assume that neurons linearly filter incoming stimuli through their receptive fields, but artificial stimuli, such as reversing gratings, often reveal nonlinear spatial processing. We investigated whether such nonlinear processing is relevant for the encoding of natural images in ganglion cells of the mouse retina. We found that standard linear receptive field models fail to capture the spiking activity for a large proportion of cells. These cells displayed pronounced sensitivity to fine spatial contrast, and local signal rectification was identified as the dominant nonlinearity. In addition, we also observed a class of nonlinear ganglion cells with opposite tuning for spatial contrast and a particular sensitivity for spatially homogeneous stimuli. Our work highlights receptive field nonlinearities as a crucial component for understanding early sensory encoding in the context of natural stimuli.

## Introduction

The natural visual world is communicated to the brain through an array of functionally distinct, parallel channels that originate in the retina (Baden et al., 2016; Roska and Meister, 2014). A classical view of retinal function advocates that the retinal output channels, represented by types of retinal ganglion cells, serve as linear filters for natural visual inputs (Atick and Redlich, 1990; Shapley, 2009). This linear picture entails that ganglion cells signal the (weighted) average of the light intensity in their center-surround spatiotemporal receptive fields (RFs) and that different ganglion cell types are discerned through differences in linear properties, such as their temporal summation profile or RF size. Support for the linear picture comes from quantitative models with linear spatio-temporal RFs that have been successful in predicting ganglion cell responses to artificial stimuli, at least when stimuli had no or coarse spatial structure at the RF scale (Aljadeff et al., 2016; Fairhall et al., 2006; Latimer et al., 2019; Ozuysal and Baccus, 2012; Pillow et al., 2008).

Some ganglion cell types, however, can carry complex nonlinear information (Gollisch and Meister, 2010), such as the direction of motion (Wei, 2018). Modeling such nonlinear responses usually requires that ganglion cells can detect spatial structure finer than the RF center, as in the case of distinguishing object from background motion (Baccus et al., 2008; Olveczky et al., 2003; Zhang et al., 2012). Sensitivity to such fine spatial structure is inconsistent with a linear RF, but has been experimentally demonstrated with artificial stimuli, such as contrast-reversing gratings, in multiple species (Demb et al., 1999; Enroth-Cugell and Robson, 1966; Krieger et al., 2017; Petrusca et al., 2007). Furthermore, linear RF models fail to predict responses of some ganglion cells to finely structured white-noise stimulation (Freeman et al., 2015; Liu et al., 2017). Such failures are linked to the nonlinear integration of excitatory signals in the RF center, which originate from presynaptic bipolar cells (Borghuis et al., 2013; Demb et al., 2001a; Turner and Rieke, 2016).

This raises the question to what extent nonlinear encoding plays a role in natural vision. In terms of spatial structure, natural stimuli lie in between finely detailed and coarse stimuli, because the light intensities of nearby regions in natural images are correlated (Burton and Moorhead, 1987), yet object boundaries can lead to edges and strong changes of stimulus intensity over short distances (Turiel and Parga, 2000). Despite early emphasis on the benefits of considering natural stimuli (Carandini et al., 2005; Felsen and Dan, 2005), only few studies have focused on whether the linear RF is a good abstraction of retinal ganglion cells during natural stimulation, and reported findings are mixed. Some studies support the idea that linear RFs are consistent with natural stimulus encoding in mouse and primate retina (Bomash et al., 2013; Nirenberg and Pandarinath, 2012), whereas others indicate that linear RF models are not generally sufficient for predicting natural scene responses in primate (Freeman et al., 2015; Heitman et al., 2016; Shah et al., 2020; Turner and Rieke, 2016) and salamander retina (Liu et al., 2017; McIntosh et al., 2017) and that their performance is cell-type specific (Turner and Rieke, 2016).

In this work, we establish a connection of spatial RF nonlinearities to natural stimulus encoding in retinal ganglion cells. We do so in the mouse retina, in which spatial integration – as measured with artificial stimuli – appears to display a broad scope (Carcieri et al., 2003), with spatially linear (Johnson et al., 2018; Krieger et al., 2017) as well as strongly nonlinear cells (Jacoby and Schwartz, 2017; Mani and Schwartz, 2017; Zhang et al., 2012). We first show that linear RF models successfully predict responses to natural images for some ganglion cells and substantially fail for others. We then connect model failure to spatial nonlinearities in the RF center. Finally, we analyze the spatial integration characteristics of specific functional cell types, showing that cells of the same type have similar properties.

## Results

### Performance of linear-nonlinear models for predicting responses to natural images varies strongly among retinal ganglion cells

In order to survey whether linear RF models could capture mouse retinal ganglion cell responses to natural images, we recorded the spiking activity of several hundred cells with multielectrode arrays. To focus on spatial aspects of natural stimulus encoding, we applied flashed, achromatic natural images, collected from three different databases (Arbeláez et al., 2011; van Hateren and van der Schaaf, 1998; Olmos and Kingdom, 2004). Images were presented for 200 ms each, separated by 800 ms of background illumination (Figure 1A). To analyze ganglion cell responses in relation to the signal inside the RF, we determined the RFs (including center and surround) from responses to spatiotemporal white noise (Figure 1B). Different cells sampled different parts of the images and displayed a variety of response patterns (Figure 1C), with apparent sensitivity to positive or negative Weber contrast and some images inducing no response (Figure 1C, top-middle). Some ganglion cells responded to both stimulus onset and offset (Figure 1D, left and middle), which may indicate ON-OFF-type RFs (Jacoby and Schwartz, 2017) or spatially nonlinear RFs (Mani and Schwartz, 2017). Furthermore, we observed both transient and sustained responses as well as response suppression (Figure 1C, bottom-right).

**Figure 1.**
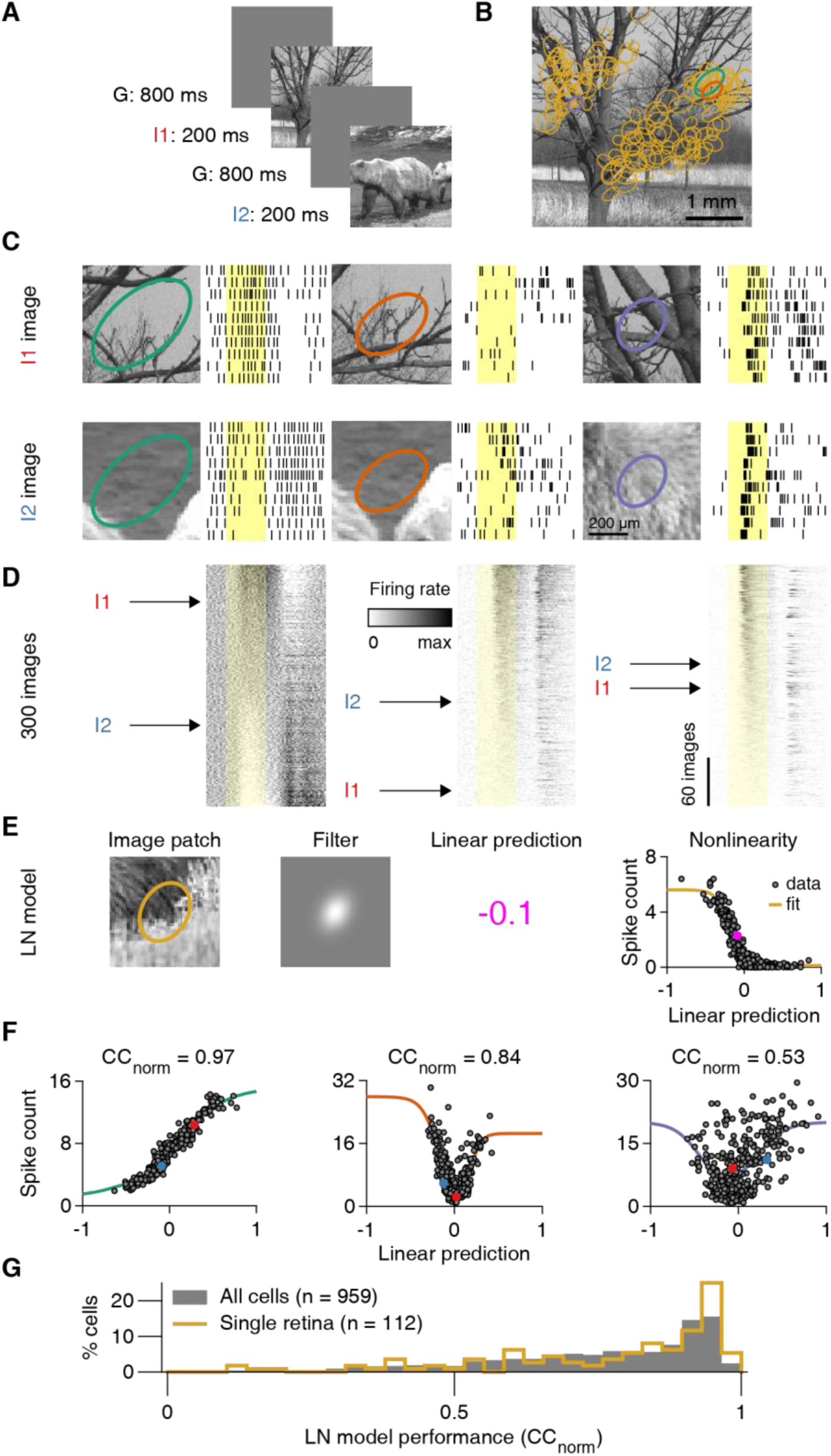
A spatially linear receptive field model often fails to predict natural image responses of retinal ganglion cells. **A)** Natural images were presented to the retina in a pseudorandom sequence for 200 ms with an inter-stimulus interval of 800 ms. **B)** Sample natural image (I1). Overlaid ellipses (light orange) represent the outlines of receptive field (RF) centers (center parts of difference-of-Gaussians fits) of 130 retinal ganglion cells from a single recording. The midline is RF-free because it contained the optic disk region. **C)** Top: Raster plots with responses of three different ganglion cells to ten presentations of image I1. Different RF outline colors correspond to different cells, also highlighted in (B). Bottom: Same as top, but for presentations of another image (I2). Yellow-shaded areas correspond to the 200-ms image presentations. **D)** PSTHs for 300 natural images, aligned to the raster plots of (C), and sorted by the average spike count during stimulus presentation. Rows corresponding to images I1 and I2 image are marked. **E)** The structure of a linear-nonlinear (LN) model that we used to predict average spike counts from natural images. The linear prediction is the inner product of the contrast values in the image patch and the filter. **F)** Spiking nonlinearities fitted to observed spike counts for the three sample cells of (C). Data points for images I1 and I2 are highlighted. The obtained normalized correlation coefficients (CC_norm_) are presented on top. **G)** LN model performance distribution for ganglion cells in a single retina preparation (light orange, same as B), and for all recorded cells (grey) from 13 preparations (9 animals).

To test whether these diverse ganglion cell responses could originate from a spatially linear RF, we measured how well a simple linear RF model could reproduce such responses. To do so, we quantified a cell’s response for each image by the average spike count following image onset. We then aimed at predicting this spike count with a linear-nonlinear (LN) model (Figure 1E). The model’s first stage is a linear spatial filter, which was estimated from a parametric fit to the spike-triggered average under white-noise stimulation. The filter captured the location, size, shape, and relative surround contribution of the spatial RF. Applying the filter to the pixel-wise Weber contrast values of a given image yielded a *linear prediction*: a single number that corresponded to the image’s net Weber contrast as seen through the cell’s RF. It quantifies how much the mean light level over the RF changed between background illumination and image presentation. The model then predicted the average spike count to the image by transforming this linear prediction with a parameterized nonlinear function, the model’s *nonlinearity*. The nonlinearity was obtained by selecting a number of images (training set) and fitting a generic function to the relation between the linear predictions and the measured responses. The obtained nonlinearity was then used to compare predictions with actual responses for the remaining images (test set), using cross-validation to quantify prediction accuracy.

For cells that linearly integrate over space, the linear prediction of the LN model should be tightly coupled to the response strength, and the relationship between the two is effectively a contrast-response function. We found cells, for example, for which the linear predictions displayed a clear, monotonic relationship to the responses, such as in Figure 1E (right, Spearman’s ρ = −0.88, n = 300 images) or 1F (left, Spearman’s ρ = 0.97, n = 300 images). An increasing monotonic relationship indicates a contrast-response function of an ON-type ganglion cell (Figure 1F), whereas a decreasing one indicates a contrast-response function of an OFF-type ganglion cell (Figure 1E, right). For such monotonic relationships, simple logistic nonlinearities provided good fits. Yet, we also found cells with a U-shaped relationship between linear predictions and responses (Figure 1F, middle). To also capture such a non-monotonic contrast-response function shape, we applied a bi-logistic nonlinearity, fitted to the contrast-response function of each cell. The bi-logistic functions captured non-monotonic nonlinearities by combining an increasing and a decreasing logistic function, but also worked well for monotonic contrast-response relations, as the weight of one logistic component naturally assumed a value near zero in the fit. Non-monotonic contrast-response functions are expected to occur in the retina for ON-OFF (Burkhardt et al., 1997) or suppressed-by-contrast ganglion cells (Jacoby et al., 2015; Tien et al., 2015), and we indeed observed both cases as indicated by U- and bell-shaped functions, respectively (Figure S1). Finally, we found cells with no apparent relationship between linear predictions and responses (Figure 1F, right and Figure 2A). For such cells, fits were poor because of the spread of data points, indicating that the LN model failed to predict responses to natural images.

**Figure 2.**
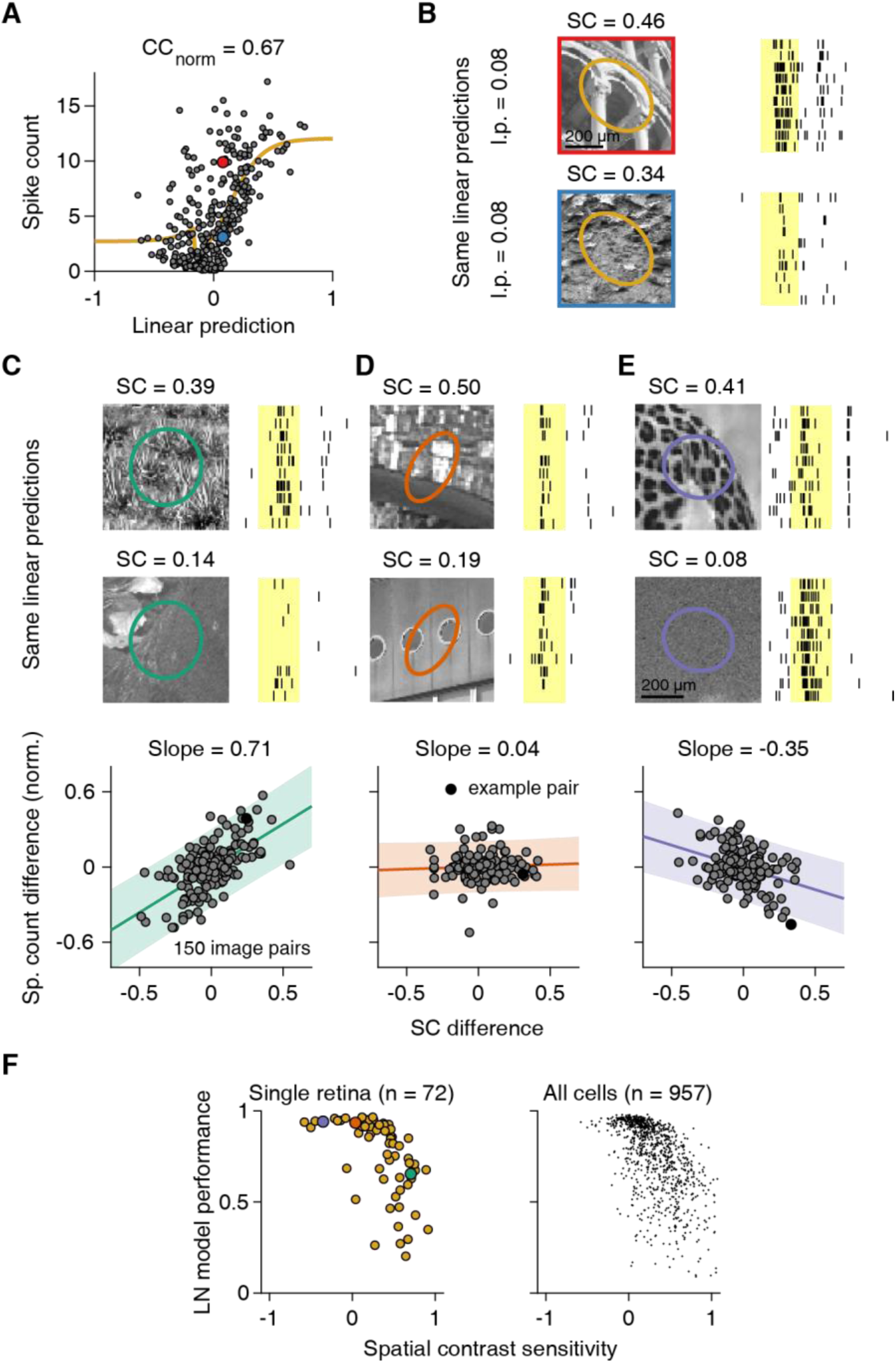
Sensitivity to natural spatial contrast is correlated with LN model performance. **A)** Output nonlinearity fit for a cell with low LN model performance. **B)** Different responses to natural images with same linear predictions (l.p.), but different spatial contrast (SC) in the receptive field (RF) center. **C)** Top: Raster plots of a sample cell’s responses to natural images with high (upper image) and low (lower image) spatial contrast in the RF center. Bottom: Relation of spatial contrast differences to average spike count differences for 150 pairs of natural images with similar generator signals in the RF center. Count differences are normalized to the maximum observed average spike count. Line and slope value correspond to least-squares estimate and shaded area to 95% confidence interval. **D-E)** Similar to (C), for two other sample cells. Shaded yellow areas in (B-E) mark the 200-ms image presentations. **F)** Relation of LN model performance to spatial contrast sensitivity, defined as the slope of the relation between spike-count differences and spatial contrast differences, as in (C-E), for ganglion cells in a single preparation (left, Spearman’s ρ = −0.73, p < 10^−100^) and for all recorded cells (right; 13 retinas, 9 animals). Data points for the sample cells in (C-E) are highlighted in the corresponding colors.

How well the LN model captures the responses can be visually assessed by how tightly the data points cluster around the fitted nonlinearities (Figure 1F), which could be quantified by how strongly prediction and response are correlated. However, part of the deviation from the fit could result from noise in the response measure, as only ten trials per image were available, rather than from an actual failure of the model. Thus, to quantify performance of the LN model, we computed a normalized correlation between response prediction and measured response, CC_norm_ (Schoppe et al., 2016), which takes the variability of responses across trials into account. Additionally, we used cross-validation by averaging CC_norm_ over ten different sets of held-out images not used to fit the nonlinearity.

Model performance varied considerably between cells. A sizeable proportion showed good model performance, indicated by a peak close to unity in the distribution of CC_norm_ values (Figure 1G). On the other hand, we observed a broad tail of cells with low CC_norm_ values, indicating different degrees of model failure, both for individual retina pieces as well as for the entire population of recorded cells. Given the variability-adjusted measure of model performance and the flexibility of the applied nonlinearity, we hypothesized that the nonlinear part of the LN model was not the source of the observed diversity in natural image encoding. We therefore focused on investigating the relation between model performance and spatial signal integration.

### Linear RF model performance correlates with spatial contrast sensitivity in the RF center

Figure 2A displays measured spike counts versus model predictions for a sample cell with low model performance. The model failure is apparent from the fact that the cell elicited widely different spike counts for images that yielded similar linear predictions of the model, corresponding to similar net contrast over the RF, and thus similar spike count predictions. The two images shown in Figure 2B, for example, had nearly identical linear predictions for the sample cell, but the cell clearly responded differently to the two images. These two images strikingly differed in their spatial structures inside the cell’s RF center (Figure 2B). We therefore quantified the spatial structure of each image within the center of a cell’s RF by computing the “spatial contrast” (see Methods), which measures the variability of image pixels inside the RF center.

To quantify the impact of spatial contrast on the spike output for a given cell, we grouped the images into pairs of similar linear predictions by the cell’s LN model. This allowed us to relate differences in spike count within a pair to differences in spatial contrast, while minimizing confounding effects of mean light-level changes inside the RF. The analysis revealed that spatial contrast was systematically related to spike count for many cells, with more spikes elicited when spatial contrast was larger (Figure 2C). Indeed, for the majority of cells (72%, n = 685/957 recorded cells), differences in spatial contrast and spike count were positively correlated, indicating that spatial contrast had a response-boosting effect beyond mean light level and that spatial integration was nonlinear.

Other cells (24%, 225/957) appeared insensitive to spatial contrast, as indicated by an approximately flat relationship between differences in spatial contrast and spike count and no significant correlation (Figure 2D). This was expected as the LN model, which is based solely on mean light level in the RF, did provide an accurate description of spike counts for some retinal ganglion cells. Unexpectedly, however, we also found a small subset of cells (5%, 47/957) that showed smaller spike counts for images with higher spatial contrast (Figure 2E). Such inverse sensitivity to spatial contrast represents a different form of nonlinear spatial integration than the response-boosting effect of spatial contrast in the majority of cells and may be described as sensitivity to spatially homogeneous stimulation.

In order to assess whether sensitivity to spatial contrast was systematically related to LN model performance, we quantified the “spatial contrast sensitivity” of a given cell by the slope of the regression line between spatial contrast and response differences, normalized by the cell’s maximum response. We found that spatial contrast sensitivity was indeed negatively correlated with LN model performance in individual experiments (e.g. Figure 2F, left; median Spearman’s ρ = −0.61, 10/13 had p < 0.05) as well as in the pooled data (Figure 2F, right; Spearman’s ρ = −0.60, p < 10^−95^, n = 957 cells). Cells for which spatial contrast boosted activity (corresponding to large positive values of spatial contrast sensitivity) were generally not as well described by the LN model. Note that the few cells with a suppressive effect of spatial contrast (negative spatial contrast sensitivity) showed fairly good LN model performance, despite the observed deviation from linear spatial integration.

### Sensitivity to fine spatial gratings alone does not predict LN model performance

Sensitivity to spatial structure on a sub-RF scale is characteristic for nonlinear RFs. A classical test for nonlinear spatial integration is to stimulate the retina with full-field contrast-reversing gratings at different spatial scales and phases (Demb et al., 1999; Hochstein and Shapley, 1976). Applying such stimuli in our recordings, we found retinal ganglion cells that clearly responded to the reversals of fine (30 µm bar width) gratings (e.g. Figure 3A-B), revealing nonlinear spatial integration under reversing gratings, similar to previous measurements in mouse retina with single-cell recordings (Krieger et al., 2017; Schwartz et al., 2012; Tien et al., 2015). These cells also responded to coarser gratings that split their RF centers in two halves. Other cells, however, barely responded to reversals of gratings for phases with zero net contrast across the RF, such as the sample cell of Figure 3C.

**Figure 3.**
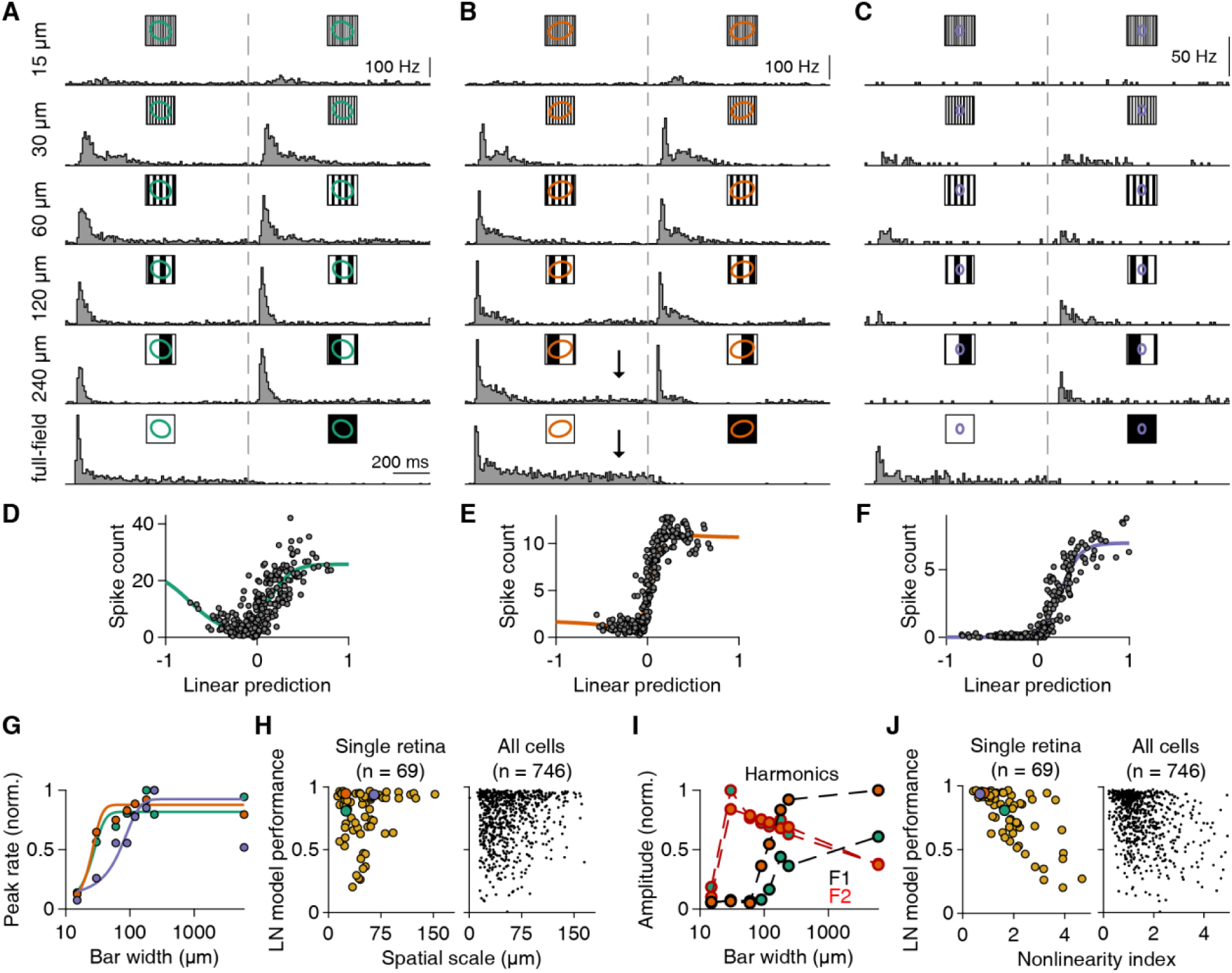
Relation of responses under reversing gratings to LN model performance. **A-C)** PSTHs of three ganglion cells to full-field contrast-reversing gratings of six different bar widths. For wider bars, the gratings were presented for multiple spatial phases, and the displayed PSTHs represent the phase with the smallest net brightness changes averaged over the receptive field (RF). Dashed gray lines correspond to the time of contrast reversal. Arrows in (B) depict the sustained component of the responses. **D-F)** Relationship between linear prediction and average spike count of the same three cells for 300 natural images. Solid lines are fitted nonlinearities. **G)** Relation of bar width to normalized peak firing rates (across time and spatial phases) for the cells in (A-C). Colors correspond to the RF colors of (A-C). **H)** Relation of LN model performance to the spatial scale of each cell for a single retinal preparation (left, Spearman’s ρ = 0.01, p = 0.94) and the total population (right) from 12 retinas, 9 animals. Cells from (A-C) are highlighted. **I)** Normalized F1 and F2 amplitudes of response harmonics across spatial frequencies for the cells of (A) and (B). **J)** Relation of LN model performance to the nonlinearity index of each cell for a single retinal preparation (left, Spearman’s ρ = - 0.7, p < 10^−10^) and the total population (right) from 12 retinas, 9 animals. The nonlinearity index is defined as the maximum F2 (across spatial frequencies) over the maximum F1 (across spatial frequencies) amplitude. Cells from (A-C) are highlighted.

Interestingly, sensitivity to contrast reversals of gratings was often unrelated to LN model performance for natural images. One of the sample cells with clear responses to fine-scale gratings (Figure 3A) had poor LN model performance for natural images (Figure 3D), whereas the other (Figure 3B) showed good model performance (Figure 3E). For the third sample cell (Figure 3C), model performance was good (Figure 3F), consistent with the observed insensitivity to grating reversals, which suggests linear spatial integration. In order to systematically compare the sensitivity to reversals of fine gratings with the LN model performance across multiple ganglion cells, we first estimated a cell’s spatial scale for detecting grating reversals by fitting a logistic curve to the cell’s peak firing rates across grating bar widths and extracting the curve’s midpoint (Figure 3G). However, there was rarely a correlation between this spatial scale and LN model performance in individual experiments (median Spearman’s ρ = 0.16, 2/12 experiments had p < 0.01; example in Figure 3H, left), and for the entire dataset, this correlation was weak, albeit significant (Spearman’s ρ = 0.12, p = 0.001, n = 746; Figure 3H, right). Thus, sensitivity to reversals of high spatial frequency gratings, typically taken as a sign for nonlinear spatial integration, does not generally imply failure of the LN model. In fact, many cells with small spatial scales showed remarkably good model performance as illustrated by the example in Figure 3E.

Examining such cells with good LN model performance but small spatial scale, we noticed differences in the firing rate profiles between different gratings. Though initial response peaks might be similar, responses could be more sustained with overall higher spike count when net-coverage of the RF with preferred contrast was larger (see arrows in Figure 3B). This response difference may be explained by spatial integration that has both a linear and a nonlinear component (e.g. resulting from partial rectification of spatially local inputs). The nonlinear component may be sufficient to generate responses to fine-scale gratings and a small measure of spatial scale, characteristic of nonlinear spatial integration. At the same time, the linear component may dominate the overall spike count under natural images, thus yielding a relatively good LN model performance. To test our hypothesis, we applied a measure that is sensitive to the spike count under reversing gratings by comparing the response Fourier components for the stimulus frequency (F1) and for twice that frequency (F2, frequency-doubled component, corresponding to responses for both reversal directions). Large F2 amplitudes, as compared to F1, are indicative of nonlinear spatial-integration effects (Hochstein and Shapley, 1976) at the level of spike counts. For the two spatially nonlinear cells of Figure 3A-B, for example, the F2 peak at 30 μm was similar, but the cell with low LN model performance had an overall smaller F1 component (Figure 3I), indicating that nonlinear effects were relatively stronger for this cell as compared to linear RF contributions.

We found that the relative amplitudes of the F1 and F2 components could indeed predict model performance better than the spatial scale. To show this, we computed a nonlinearity index as the ratio of the maximal F2 amplitude (over grating widths and phases) and the maximal F1 amplitude (see Methods). The nonlinearity index was negatively correlated to LN model performance under natural images both in single experiments (median Spearman’s ρ = −0.40, 9/12 experiments had p < 0.01) as well as in the whole population (Spearman’s ρ = −0.35, p < 10^−22^, n = 746). Thus, the relative degree of nonlinear spatial integration appears to play a role in determining ganglion cell responses to natural images.

Despite the correlation between the nonlinearity index with LN model performance, the classical analysis with contrast-reversing gratings had some drawbacks. Firstly, for ON-OFF cells, the approach cannot distinguish between nonlinear integration over space or over ON-type versus OFF-type inputs, as both phenomena can lead to large F2 components. Secondly, the analysis only detects that some rectification of non-preferred contrasts exists (as effects of preferred and non-preferred contrast do not cancel out), but it does not offer information about how preferred contrast signals at different locations inside the RF are combined.

### Responses to contrast combinations inside the RF reveal the components of natural spatial contrast sensitivity

To overcome the shortcomings of classical contrast-reversing grating stimulation and explore the relationship between spatial contrast sensitivity and LN model performance more systematically, we designed a checkerboard flash stimulus that tests a range of contrast combinations. This stimulus was based on the idea of independently stimulating two separate sets of spatial subunits within a cell’s RF with different inputs (Bölinger and Gollisch, 2012; Takeshita and Gollisch, 2014). Concretely, we flashed a batch of varied checkerboards onto the retina (Figure 4A, top). The contrasts of the two sets of tiles, or spatial inputs, were sampled from the stimulus space of pairs of contrast values (Figure 4A, bottom-right) to systematically explore a wide range of contrast combinations. To directly compare responses between artificial and natural stimuli, we flashed the contrast pairs for 200 ms each (the same duration as for the natural images) in a pseudorandom sequence, collecting four to five trials per pair. The subfields of the checkerboard spanned 105 μm to the side, approximately half of the average mouse RF center, to provide a strong, yet spatially structured stimulus inside the RF.

**Figure 4.**
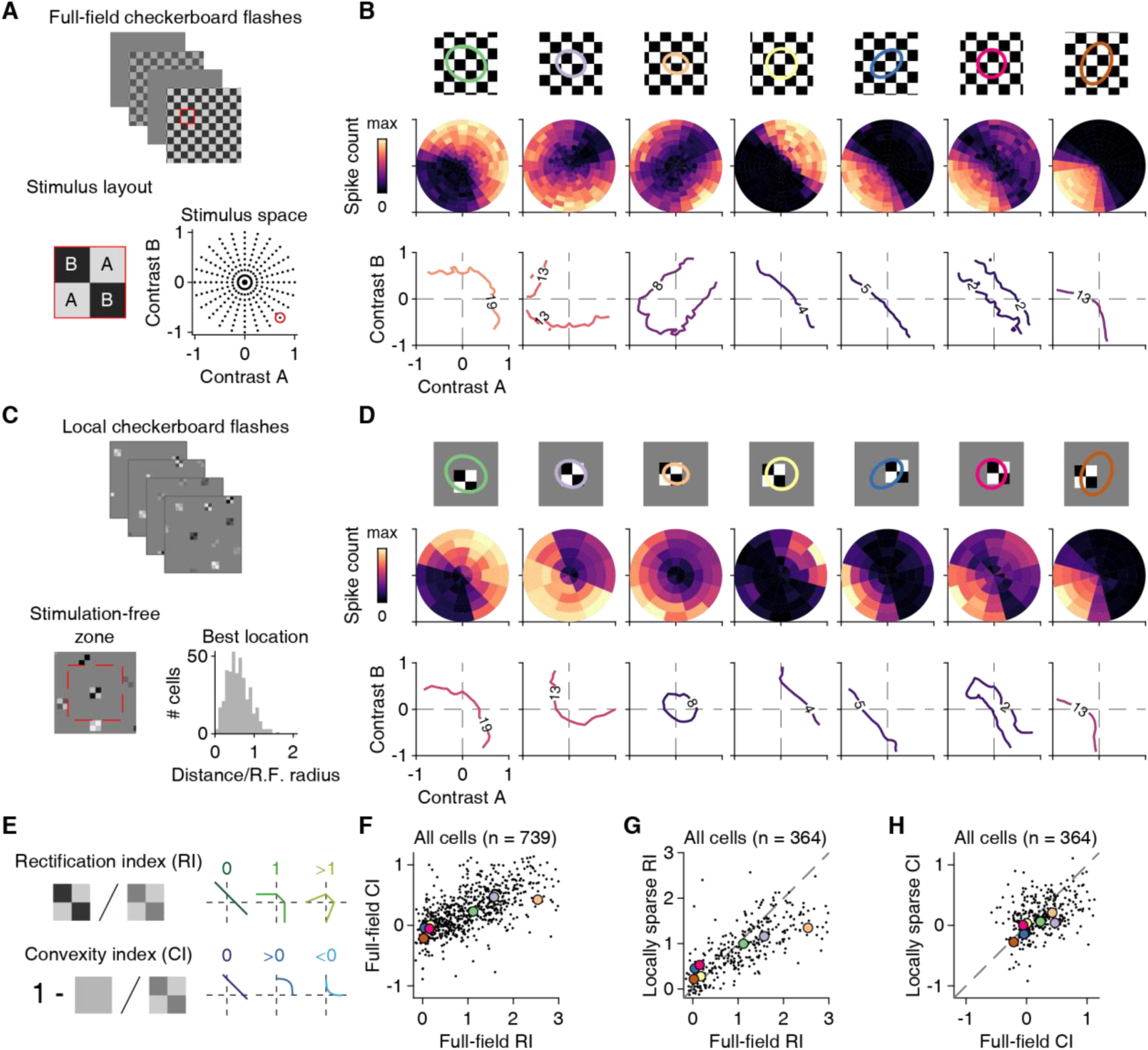
Stimulation with contrast combinations reveals nonlinearities in spatial input integration. **A)** Depiction of the applied stimulus space, comprising flashed checkerboards with contrast combinations (A,B) sampled along different directions in stimulus space. The dots in the stimulus space on the lower right mark all contrast combinations applied in the experiment. The contrast combination identified by the red circle corresponds to the example on the left, which shows a 2×2 cutout from the stimulus frame shown on top (marked by the red square). **B)** Top: Receptive field (RF) center outlines of seven sample ganglion cells, relative to the stimulus layout, with tiles for contrast A and B shown in white and black, respectively. Middle: Color-coded average spike counts for all tested contrast pairs in the stimulus space. Bottom: Iso-response contours in the stimulus space for a selected spike count, indicated by the number on the contour and the contour’s color. **C)** Top: Example frames of the locally sparse stimulus. Bottom: Display of the region (dashed red line) around a stimulus location that was excluded from further selection of stimulus locations (left). Distribution of RF center distance to the center point of the closest grid square (n = 404), normalized by the RF radius. **D)** Same as (B), but for the locally sparse stimulus. **E)** Schematic depiction of how rectification and convexity indices are computed and how they are related to different shapes of contour lines. **F)** Relation of full-field rectification and convexity indices in the pooled ganglion cell data from 13 retinas, 9 animals. **G)** Relation of full-field and locally sparse rectification indices in the pooled ganglion cell data from 6 retinas, 4 animals. Dashed line is the equality line. **H)** Same as (G), but for the convexity index.

The checkerboard flashes revealed a variety of spatial integration profiles among different retinal ganglion cells. To extract these profiles, we visualized the responses (defined as the average spike count over 250 ms after stimulus onset, equivalent to the response measure under natural images) as color maps over the stimulus space of contrast pairs (Figure 4B, middle row). We then calculated iso-response contour lines (Figure 4B, bottom row), which trace out those contrast pairs that led to the same response (here number of spikes). The shape of the contour lines is indicative of the type of subunit nonlinearity (Bölinger and Gollisch, 2012; Gollisch and Herz, 2012; Maheswaranathan et al., 2018). Straight contour lines, for example, correspond to linear integration of the two inputs, whereas curved ones reflect the existence of a nonlinearity and indicate the type of nonlinearity in the shape of the contour.

We found both ON and OFF varieties of nonlinear cells (Figure 4B, Cells 1-2), with contour lines curving away from the origin. This nonlinear signature may result from an expansive transformation of local signals, such as by a threshold-quadratic functions (Bölinger and Gollisch, 2012), or by a sigmoid with high threshold (Maheswaranathan et al., 2018). We also found linear ON and OFF ganglion cells, with straight contour lines (Figure 4B, Cells 4-5). Furthermore, our approach allowed us to visualize the spatial integration profiles of ON-OFF cells, and distinguish between spatially nonlinear and linear ON-OFF cells (Figure 4B, Cells 3 and 6). Linear ON-OFF cells responded mostly to net-increases or decreases of light intensity, but not when the two contrast signals cancelled each other, leading to parallel contour lines (Cell 6). On the other hand, nonlinear ON-OFF cells often had closed or nearly closed contour lines, corresponding to strong responses also for contrast combinations with opposing signs (Cell 3). Finally, we identified a unique nonlinear spatial integration profile in some cells, characterized by contour lines curving towards the origin (Figure 4B, Cell 7). Such a profile indicates a particular preference to a spatially homogeneous change in light level, as has been previously observed in the salamander retina (Bölinger and Gollisch, 2012). We mainly found such profiles for OFF-type ganglion cells, but occasionally in ON-type cells as well (6/27 cells were ON-type).

For comparison, we also devised a local version of checkerboard flashes in order to assess potential contributions of the RF surround to nonlinear spatial integration. Here, the display of each contrast combination was spatially restricted to a patch of 2×2 tiles of the checkerboard, which roughly corresponds to typical RF center sizes. To nonetheless cover the entire recording area and obtain sufficient sampling of contrast combinations, multiple randomly chosen patches, obeying local sparsity (Hawrylycz et al., 2016; de Vries et al., 2018), were displayed simultaneously (Figure 4C), and fewer contrast combinations were sampled as compared to the full-field version of the stimulus. For further analysis, we selected for each cell the patch location closest to the RF center. This generally lay not further away than one RF radius (Figure 4C, bottom right), indicating good overlap of the analyzed patch location with the RF center. We found that spatial integration profiles, as captured by the shape of the contour lines in stimulus space, were qualitatively similar under local stimulation as compared to full-field stimulation (Figure 4D).

For both full-field and local stimulation, the examples show that the contour lines can deviate from straight lines in different ways. Rectification of non-preferred inputs, for example, becomes visible by how the contour line bends as it progresses from the quadrant in stimulus space that corresponds to preferred contrast for both stimulus components (upper right quadrant for ON cells; lower left for OFF cells) to the two neighboring quadrants that combine positive and negative contrast. In addition, there is also nonlinear integration of preferred contrast, which is visible in a nonlinear shape of the contour line inside the quadrant that corresponds to preferred contrast of both stimulus components. To quantify these nonlinear contributions, we devised two corresponding indices (Figure 4E). We calculated a rectification index by comparing responses to flashes where both components had opposing, equal-magnitude contrast with responses when only a single stimulus component was used (Molnar et al., 2009). Full rectification leads to equal responses for both configurations and an index of unity, whereas linear integration would make the opposing-contrast configuration effectively a null stimulus, resulting in no response and an index of zero. Similarly, we computed a convexity index by comparing responses from using just one spatial input at a specific contrast level with responses from using both inputs at half that contrast level. A convexity index of zero corresponds to linearity (equal responses for a single component at full contrast and for two components at half contrast), whereas values smaller or larger than zero correspond to increased or decreased preference for homogeneous stimuli, respectively (see Figure S2 for sample responses used to compute the indices). Over the population of all recorded cells, the two indices were correlated (Figure 4F for the full-field indices; Spearman’s ρ = 0.68, p < 10^−100^, n = 739), indicating that the two nonlinearity components often coexist.

To systematically compare full-field and local spatial integration profiles, we compared rectification and convexity indices across the two conditions. Although both indices displayed a significant change between local and full-field stimulation (Wilcoxon signed-rank test, p < 10^−10^ for the rectification and p < 10^−6^ for the convexity index), the values were correlated between the two conditions (Spearman’s ρ = 0.80, p < 10^−73^ for the rectification and Spearman’s ρ = 0.39, p < 10^−12^ for the convexity index), indicating that cells retained their relative characteristics of nonlinear spatial integration, in particular regarding rectification (Figure 4G). One subtle change was that for many cells with convexity index >0 in the full-field stimulus, the index became ∼0 for the local stimulus (Figure 4H), corresponding to a less outward-bulging shape of the contour line in the quadrant of preferred contrast (visible in the first two examples when comparing Figures 4B and 4D). This indicates that homogeneous stimulation was less effective under full-field conditions (relatively larger contrast values are needed to reach the activation level of the contour line). A likely reason is that spatially homogeneous stimuli cause relatively stronger surround suppression than spatially structured stimuli, which could arise from linear spatial integration in the surround.

How are the extracted components of nonlinear spatial integration related to responses under natural images? We found that spatial contrast sensitivity, as determined from the responses to natural images (cf. Figure 2D), was correlated with both the rectification (Spearman’s ρ = 0.71, p < 10^−112^) and the convexity index (Spearman’s ρ = 0.58, p < 10^−67^), as obtained from full-field stimulation (Figure 5A and 5C, top). Similar results were also found for the indices obtained from local stimulation (Figure 5B and 5D, top; Spearman’s ρ = 0.70, p < 10^−53^ for rectification and ρ = 0.33, p < 10^−9^ for convexity). At the same time, the rectification indices from both full-field (Spearman’s ρ = −0.71, p < 10^−113^) and local stimulation (Spearman’s ρ = −0.69, p < 10^−51^) were good predictors of LN model performance for natural images (Figure 5A-B, bottom), to an extent much larger than the nonlinearity indices extracted from contrast-reversing gratings. The convexity indices from full-field (Spearman’s ρ = −0.56, p < 10^−60^) and local stimulation (Spearman’s ρ = −0.40, p < 10^−14^) were also predictors of LN model performance (Figure 5C-D, bottom), but to a smaller extent than rectification indices. We thus concluded that the degree of rectification of spatial inputs in the RF center is a primary factor that shapes ganglion cell responses to natural images and determines whether responses can be captured by the LN model.

**Figure 5.**
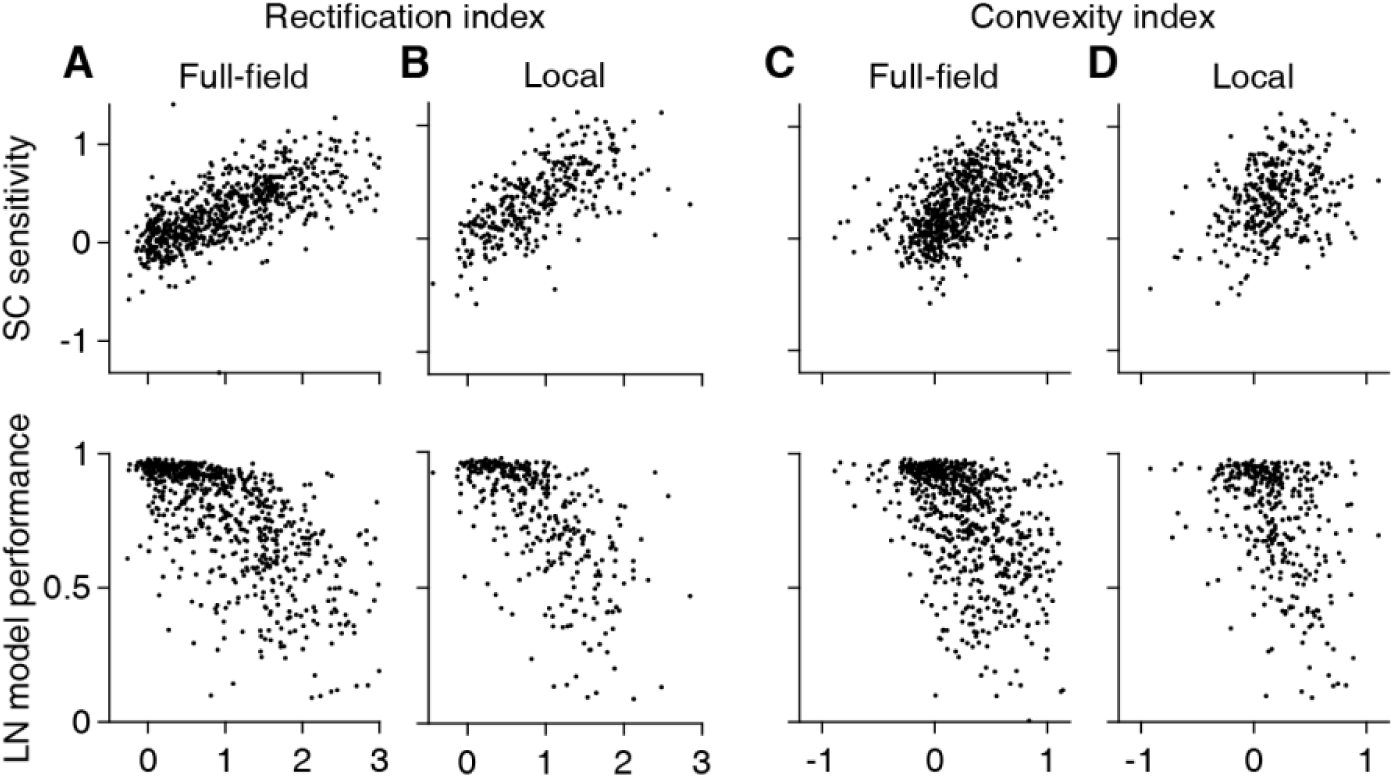
Spatial input nonlinearities curtail spatial contrast sensitivity for natural images. **A)** Relation of the spatial contrast (SC) sensitivity (top) and linear-nonlinear (LN) model performance (bottom), measured with natural images, to the full-field rectification index in the pooled cell data (n = 739) from 13 retinas, 9 animals. **B)** Same as (A), but using the data from the local checkerboard flash stimulus in the pooled cell data (n = 364) from 6 retinas, 4 animals. **C-D)** Same as (A-B), but for the convexity index.

### The spatial scale of contrast sensitivity for natural images

We next asked on what spatial scale nonlinearities are relevant for encoding natural images. To do so, we compared responses under original natural images and blurred versions (Figure 6), similar to previous analyses with white-noise patterns (Jacoby and Schwartz, 2017; Johnson et al., 2018; Mani and Schwartz, 2017; Schwartz et al., 2012). The blurring with a given spatial scale corresponds to low-pass filtering and removes fine spatial structure below this scale while keeping the mean intensity over larger regions approximately constant. At a blurring scale close to a cell’s RF center diameter, blurring should diminish spatial contrast within the RF while keeping the mean light intensity approximately unchanged. Figures 6A-C compare responses to natural images and their blurred versions for three sample cells. At a scale of 240 µm, the blurring generally reduced responses for the first cell (Figure 6A, middle; Wilcoxon signed-rank test, n = 40 images, p < 10^−6^) but left responses for the second largely unaffected (Figure 6B, middle; p = 0.18), and for the third cell even led to increased spike count (Figure 6C, middle; p = 0.02).

**Figure 6.**
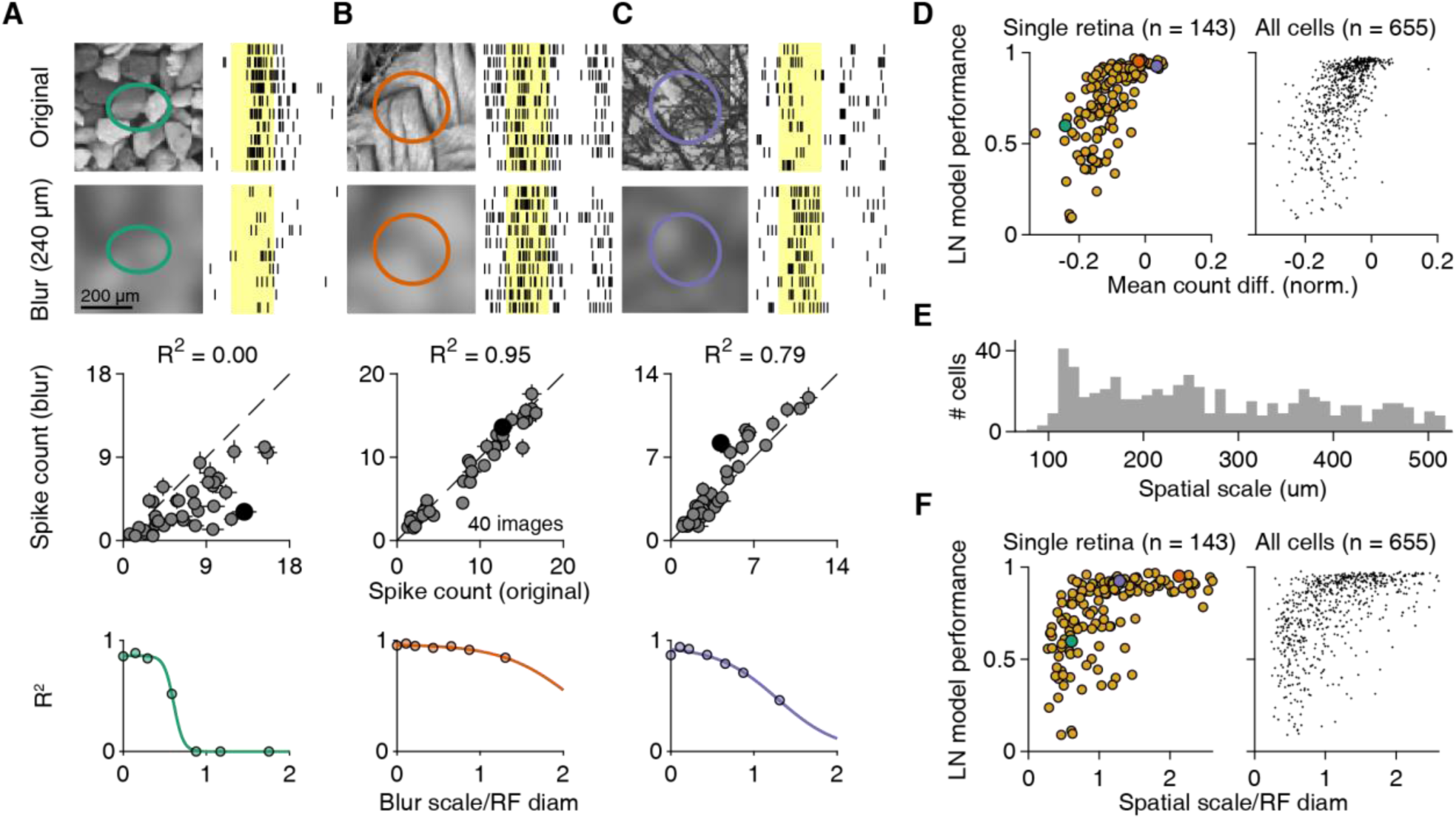
The scale of spatial contrast sensitivity for natural images. **A-C)** Top: Responses of three sample cells to original natural images and their blurred versions for ten trials. Images were blurred with a Gaussian function (scale 4σ = 240 μm). Shaded yellow areas mark the 200-ms image presentations. Middle: Relation of the average spike count for presentations of 40 natural images and their blurred counterparts for each cell of (A). Filled black dots correspond to the image pairs shown above. Error bars denote SEM. Dashed line is the equality line. Bottom: Coefficient of determination (R^2^) between original and blurred spike counts for different degrees of blurring, normalized by the RF diameter of each cell. Colored lines correspond to logistic fits. **D)** Relation between linear-nonlinear (LN) model performance for natural images and the mean (across images) spike count difference for each cell from a single retina (left, Spearman’s ρ = −0.73, p < 10^−24^) and the pooled ganglion cell population (right, Spearman’s ρ = 0.67, p < 10^−86^) from 9 retinas, 6 animals. The differences were normalized to the maximum average spike count observed for the cell. Cells from (A-C) are highlighted. **E)** The distribution of spatial scale across the pooled ganglion cell population (n = 655). Spatial scale was defined as the midpoint of the logistic functions in (A-C, bottom). **F)** Relation of spatial scale, normalized by the RF diameter, to LN model performance for natural images for a single retina (left, Spearman’s ρ = 0.61, p < 10^−15^) and the pooled ganglion cell population (right, Spearman’s ρ = 0.60, p < 10^−64^) from 9 retinas, 6 animals. Cells from (A-C) are highlighted.

To quantify the blurring effects for all cells, we calculated the mean response difference between the blurred and the original version of the images, normalized by the cell’s maximum response over all images. This spike count difference was correlated to LN model performance for natural images (Figure 6D) in both individual experiments (median Spearman’s ρ = 0.64, 9/9 experiments had p < 0.01), and in the pooled population (Spearman’s ρ = 0.67, p < 10^−86^). Similarly, the spike count difference under blurring was also negatively correlated to spatial contrast sensitivity (Figure S3) in both individual experiments (median Spearman’s ρ = −0.65, 7/9 experiments had p < 0.001), and in the pooled population (Spearman’s ρ = −0.73, p < 10^−110^). Thus, cells that were more strongly affected by the blurring generally displayed worse LN model performance and had a stronger dependence of spike count on spatial contrast. This confirms the effect of spatial structure inside the RF for determining responses to natural images in particular ganglion cells.

When analyzing responses across different blurring scales, we observed that cells sensitive to spatial contrast reduced their spike counts already at scales smaller that their RF center (Figure 6A, bottom). To quantify the spatial scale of blurring sensitivity for each cell, we measured the similarity between responses to original and blurred images by calculating the corresponding coefficient of determination (R^2^), which is unity when responses with and without blurring are identical and falls off towards zero as responses to blurred images deviate more and more from the original responses. Analogous to the analysis of contrast-reversing gratings, we fitted logistic functions to the decay of R^2^ with blurring scale and defined the spatial scale as the midpoint of the logistic function. The obtained spatial scales ranged from 100 to 500 μm (Figure 6E) and were only weakly correlated with the spatial scales measured with contrast-reversing gratings (Spearman’s ρ = 0.12, p = 0.006). And unlike the spatial scale obtained from reversing gratings, the spatial scale from blurred images (normalized by the RF center diameter) was strongly related to LN model performance (Figure 3F) in both individual experiments (median Spearman’s ρ = 0.59, 9/9 experiments had p < 0.05) and in the pooled population (Spearman’s ρ = 0.60, p < 10^−64^).

### Spatial contrast sensitivity differs among retinal ganglion cell classes

The analyses so far have shown that the characteristics of spatial integration are consistent for individual ganglion cells across different stimulus conditions, including natural and artificial stimuli. We thus hypothesized that they reflect cell-type specific properties. To test this hypothesis, we looked at three readily identifiable cell classes, detected through a standard set of artificial stimuli.

Firstly, we focused on image-recurrence-sensitive (IRS) cells, which form a single functional cell type in the mouse retina and which correspond to transient OFF-alpha ganglion cells (Krishnamoorthy et al., 2017). We identified IRS cells by their characteristic response peaks to rapid shifts of a grating with no net-displacement of the grating position (Figure 7C). As expected, IRS cells were all OFF-type, with fast temporal filters and tiling RFs. For these cells, all our spatial integration measures displayed relatively narrow distributions. LN model performance for IRS cells was high (median = 0.94, n = 29), suggesting linear spatial integration. However, rather than showing no sensitivity to spatial contrast, the distribution of spatial contrast sensitivity for IRS cells was significantly shifted towards negative values (median = −0.11, n = 29, Wilcoxon sign-rank test, p = 0.005). This indicates that IRS cells had a particular preference for spatially homogeneous natural stimuli. Specifically, about half (15/29) of the IRS cells were inversely sensitive to spatial contrast of natural images. In terms of spatial integration measured by the checkerboard flashes, most IRS cells showed profiles such as the one in Figure 4B (Cell 7), with low rectification (median = 0.12, n = 28) and slightly negative convexity indices, yet not significantly different from zero (median = −0.06, Wilcoxon sign-rank test, p = 0.07).

**Figure 7.**
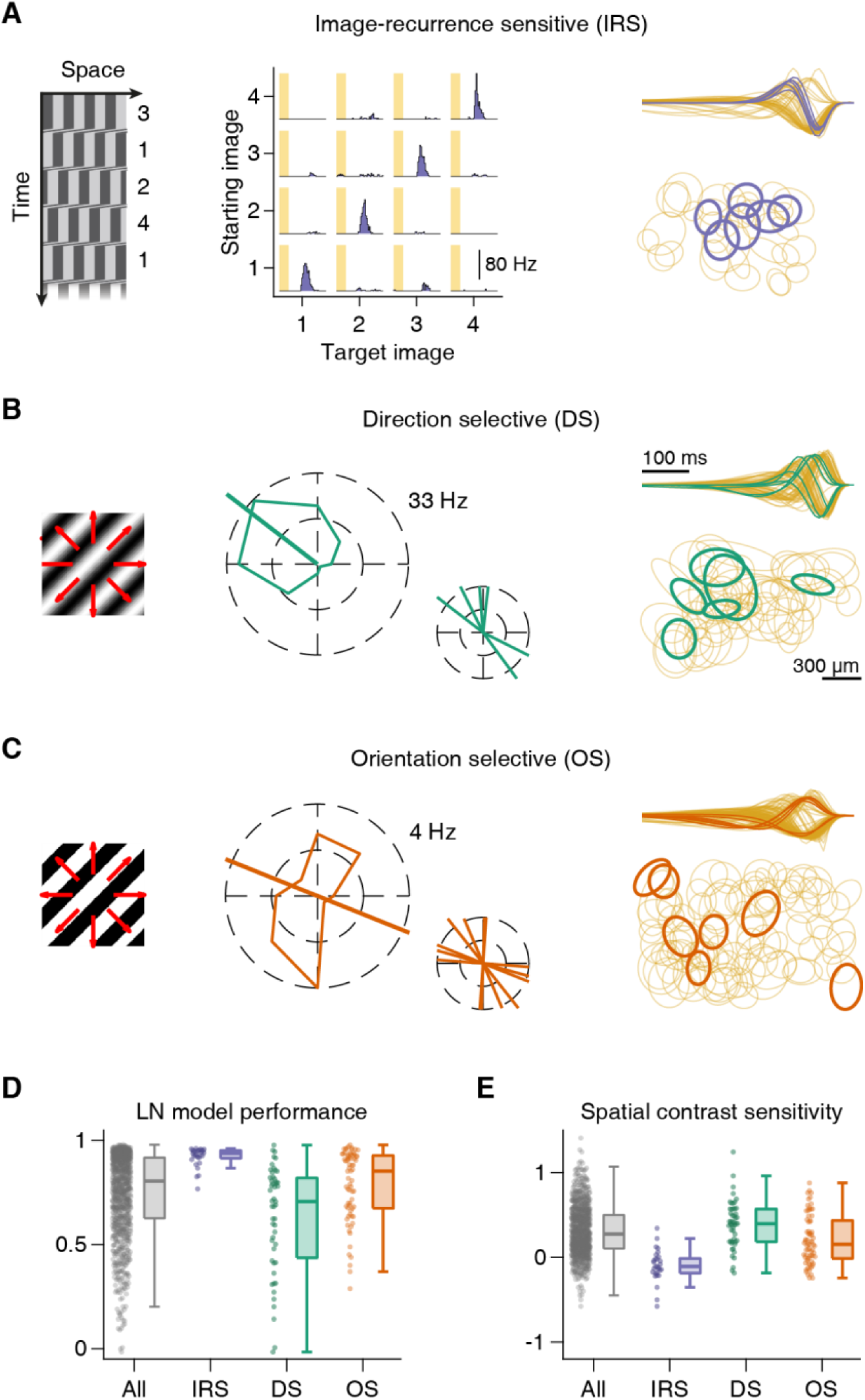
Spatial contrast integration in three different functional cell classes. **A)** Image-recurrence sensitive (IRS) cells were identified with a sequence of saccadic-like grating movements (left), shown as a space-time plot with time going downwards. Numbers denote the four different fixation positions. PSTHs of an example IRS cell for the 16 possible transitions (middle), with shaded areas marking the 100-ms transitions. Temporal filters (normalized to unit norm) of IRS cells in a single retina (right-top), overlaid on top of the temporal filters of all recorded cells. Below, the receptive field (RF) centers of IRs cells (right-bottom), overlaid on top of some of the recorded RFs. **B)** Direction selective (DS) ganglion cells were identified from their responses to slow drifting sinusoidal gratings (left). Tuning curve of a sample DS cell (middle), along with the preferred direction of all DS cells in a single recording. Temporal filters and RFs are shown as in (A). **C)** Orientation selective (OS) ganglion cells were identified with drifting square-wave gratings (left), moving at higher speeds than in (B). Tuning curve of a sample OS cell (middle), along with the preferred orientations of all OS cells in a single recording. Temporal filters and RFs are shown as in (A). **D)** Distributions of LN model performance under natural stimuli for IRS (n = 29), DS (n = 51), and OS (n = 70) cells from 13 retinas, 9 animals. Gray corresponds to all cells. **E)** Same as (D), but for spatial contrast sensitivity.

Secondly, we tested direction-selective (DS) ganglion cells (Figure 7A), detected through their responses to drifting gratings. DS cells had either ON- or OFF-type temporal filters (Figure 7A, right-top), with OFF-type filters likely corresponding to ON-OFF DS cells, as suggested by U-shaped nonlinearities in LN models obtained from white-noise stimulation (data not shown). DS cells generally displayed rather nonlinear spatial integration for natural images (Figure 7D-E), with low LN model performance (median = 0.70, n = 51) and significant spatial contrast sensitivity (median = 0.40, n = 51, Wilcoxon sign-rank test, p < 10^−7^). As expected for cells sensitive to spatial contrast, their full-field rectification indices were rather large (median = 1.10, n = 35), and their convexity indices were significantly larger than zero (median = 0.28, n = 35, Wilcoxon sign-rank test, p < 10^−3^). Thus, DS cells were rather nonlinear in their spatial integration, as seen by their responses to both natural and artificial stimuli.

Finally, using drifting gratings with higher speed, we identified orientation-selective (OS) ganglion cells (Figure 7B). We found both ON- and OFF-type OS cells, possibly corresponding to the recently described different classes in the mouse retina (Nath and Schwartz, 2016, 2017). OS cells displayed characteristics of spatial integration that lay in between IRS and DS cells. Compared to DS cells, for example, OS cells showed better (Wilcoxon rank-sum test, p < 10^−4^) LN model performance, yet many cells still revealed poor model predictions (median = 0.85, n = 70). Likewise, spatial contrast sensitivity was lower that for DS cells, yet still significantly larger than zero (median = 0.15, n = 77, Wilcoxon sign-rank test, p = 0.002). This was also reflected in the OS cells’ responses to the checkerboard flashes, with slightly lower full-field rectification (median = 0.57, n = 53) and convexity indices (median = 0.20, n = 53) as compared to DS cells, yet with both indices significantly larger than zero (Wilcoxon sign-rank test, p < 10^−9^ for rectification and p = 0.002 for convexity indices).

Examining the distributions of these measures for OS cells more closely, we observed that they actually appeared to be bimodal (Figure S4A), which may indicate that different types of OS cells differ in how nonlinear their spatial integration is. Indeed, we found that ON-type OS cells showed fairly linear spatial integration characteristics (Figure S4A), while OFF-type OS cells could be nicely clustered into two separate groups, one with linear spatial integration and good LN model performance and another with nonlinear spatial integration and poorer LN model performance (Figure S4A-B). Interestingly, the linear and nonlinear OFF OS cells also differed systematically in their preferred orientations (Figure S4C). We may thus speculate that the previously described classes of OS cells with different contrast preference or preference for horizontal or vertical orientations (Nath and Schwartz, 2016, 2017) are also separated by having either linear or nonlinear spatial integration.

## Discussion

In this work, we aimed at directly addressing the question whether retinal ganglion cell responses to natural stimuli are consistent with a linear RF. We found that this is the case only for a subset of cells in the mouse retina under natural images. With targeted artificial stimulation, we were able to link these differences in natural image encoding to different degrees of nonlinear spatial integration inside the RF center in different cell classes.

### Diversity in natural stimulus encoding among the retina’s output channels

Using a simple linear RF model, we observed multiple facets of natural image encoding in the mouse retina. We found ganglion cells that were consistent and others that were inconsistent – to different extents – with linear RFs. Out of the few previous studies with natural stimuli in the mouse retina, one supports generally linear RFs in mouse ganglion cells (Nirenberg and Pandarinath, 2012). However, both spatially linear and nonlinear ganglion cell types had been identified in the mouse retina with artificial stimuli. For example the PixON (Johnson et al., 2018) or the sustained OFF-alpha cells (Krieger et al., 2017) appear to have linear RFs, whereas nonlinear RF properties can be detected for ON-delayed or ON-OFF DS cells (Mani and Schwartz, 2017). Here, we showed that DS cells, like several other ganglion cells, are spatially nonlinear also for natural stimuli.

The degree and type of spatial nonlinearity appears to differ between retinal ganglion cell types, yet functional cell type classification schemes rely mostly on linear model components, such as RF size and temporal filter shapes (Baden et al., 2016; Chichilnisky and Kalmar, 2002; Franke et al., 2017; Jouty et al., 2018; Ravi et al., 2018; Rhoades et al., 2019). Including characteristics of nonlinear spatial integration – such as LN model performance, subunit rectification, or spatial scale of linear integration – may help to better distinguish cell types. For example, the mouse retina contains at least four subtypes of OS cells, two of which (ON- and OFF-type) are tuned to horizontal and the other two (again ON and OFF) to vertical orientations (Nath and Schwartz, 2016, 2017). Here, we found distinct groups of OS cells with linear and nonlinear spatial integration, suggesting that OS subtypes might differ not only in contrast preference or preferred orientation, but also in how they integrate spatial contrast, which provides additional information for separating and identifying subtypes of OS cells.

When considering that synaptic transmission is often inherently nonlinear, the occurrence of linear RFs may actually be surprising (Shapley, 2009). In the salamander retina, for example, nonlinear ganglion cell RF centers and surrounds seem to be the norm (Bölinger and Gollisch, 2012; Takeshita and Gollisch, 2014). Linear RFs may be a property of mammalian retinas, as they have been described also in cat, rabbit and macaque retinas (Enroth-Cugell and Robson, 1966; Molnar et al., 2009; Petrusca et al., 2007; White et al., 2002), and may have specifically evolved to provide raw information about illumination patterns to the cortex for further processing (Roska and Meister, 2014).

### A cell class with particular sensitivity to spatial homogeneity of natural images

We identified cells in the mouse retina with particular sensitivity to spatially homogeneous image parts. Specifically, these cells were inversely sensitive to spatial contrast: though well described by an LN model, they respond more strongly to homogeneous stimuli than to structured stimuli of equal light level. This feature is not to be confused with the characteristics of suppressed-by-contrast cells (Jacoby and Schwartz, 2018), which are suppressed below baseline activity by (temporal) contrast, whereas the homogeneity-preferring cells here are particularly strongly activated by homogenous spatial stimuli. This is reminiscent of the homogeneity detectors that have been described in the salamander retina (Bölinger and Gollisch, 2012), although the latter showed rectification of non-preferred contrasts, unlike the homogeneity-sensitive cells described here. Note also that these cells, through their particular sensitivity to homogeneous stimuli, could provide information about image focus; blurring through de-focusing will increase activity for this cell type and simultaneously decrease activity for spatial-contrast-sensitive cells, such as ONdelayed cells (Mani and Schwartz, 2017), which have been implicated in focus-sensing functions. A readout based on activity differences between cells of opposite tuning under image blur could provide a code for image focus that is particularly robust, for example to variations in contrast and spatial structure (Kühn and Gollisch, 2019).

IRS cells appear to be part of the homogeneity-sensitive cells. The IRS cells correspond to transient OFF-alpha cells (Krishnamoorthy et al., 2017) and should therefore also match the PV5 ganglion cells, which have been shown to be approach-sensitive (Münch et al., 2009). It seems that approach sensitivity, image-recurrence sensitivity, and sensitivity to homogeneous natural images may rely on the same circuit component: strong, local (glycinergic) ON-type inhibition, which transient OFF-alpha cells are known to receive (van Wyk et al., 2009) and which needs to be suppressed – perhaps below baseline activation level – by OFF-type stimuli for maximal activity.

### Mechanisms of nonlinear spatial integration in natural image responses

The central nonlinear operation governing spatial integration appears to be signal rectification, with different ganglion cells showing different degrees of rectification. We demonstrated this for both artificial and natural stimuli, similar to what was shown in the macaque retina for OFF parasol cells (Turner and Rieke, 2016). Signal rectification is usually attributed to the rectified excitation that bipolar cells provide to the ganglion cell RF (Demb et al., 2001b). Our measure of spatial scale that separates linear and nonlinear spatial integration in natural images yielded values of 100 μm and larger. These estimates exceeded the reported range of bipolar cell RFs, whose diameters are around 40-60 μm in the mouse retina (Berntson and Taylor, 2000; Franke et al., 2017; Schwartz et al., 2012), potentially because spatial scales are enlarged by the spatial correlations contained in natural images.

For some ganglion cells, spatial scales may also come out to be larger than individual bipolar cell RFs because of electrical coupling between bipolar cells (Kuo et al., 2016), especially for stimuli with considerable spatiotemporal correlations, such as natural stimuli. Furthermore, for some ganglion cells in the primate retina, enlarged spatial scales of stimulus integration seem to result from nonlinear dendritic processing through active conductances in the ganglion cell after some integration of bipolar cell signals has already taken place (Rhoades et al., 2019). Thus, the structural basis of spatial nonlinearities for natural stimuli are likely either individual bipolar cells or groups of nearby bipolar cells.

That the degree of nonlinear spatial integration covers a wide, continuous range rather than separating ganglion cells into distinct linear and nonlinear classes can be explained by different degrees of partial rectification in the RF. How is the degree of rectification controlled? A potential cell-intrinsic mechanism is that baseline activity of bipolar cell synapses can vary, allowing some synapses to modulate transmitter release in both directions, whereas others cannot decrease activity much below baseline and thus provide rectified transmitter release. For example, regarding inputs to Y-type ganglion cells in guinea pig and mouse, the basal glutamate release is higher at the more linear ON compared to OFF bipolar cell terminals (Borghuis et al., 2013; Zaghloul et al., 2003). Another candidate mechanism is a circuit motif: the level of crossover inhibition carried by glycinergic amacrine cells (Werblin, 2010). Bipolar cell signals alone may primarily provide rectified activation for preferred contrast, and (partial) linearization comes from crossover inhibition, which adds suppression for non-preferred contrast. The crossover inhibition can act on bipolar cell terminals (Molnar et al., 2009) or directly on the ganglion cell, and its gain, relative to the gain of the preferred-contrast excitation, would determine the degree of nonlinearity in spatial integration.

### Limitations of this study

Using multielectrode array recordings (e.g. rather than targeted single-cell recordings) has the disadvantage that clearly identifying multiple cell types is (still) difficult. On the other side, owing to the high-throughput nature of the recordings, the multielectrode array recordings can provide an overview of the diversity of responses. A further issue arises with the use of natural images to study spatial integration. Since RFs of recorded ganglion cells are distributed over the fairly broad range of the recording sites of the multielectrode array, different cells are stimulated by different regions in the images. This creates variability in the findings. For example, a nonlinear cell that happens to primarily experience image patches with little spatial structure may show reasonably good performance by the LN model. This broadens the distribution of model performance even within a single cell type.

We used flashed image presentations because our study was focused on spatial integration. Yet, this approach is insensitive to nonlinearities that become relevant when the temporal structure of stimuli is considered. For example, IRS cells, which we here reported as being rather linear for the encoding of images flashed in isolation, can reveal nonlinearities when rapid image transitions are considered, for which disinhibitory interactions mediate a sensitivity to recurring spatial patterns (Krishnamoorthy et al., 2017). Additionally, we focused on a single light level, but spatial nonlinearities may change with light level: sustained ON-alpha ganglion cells in the mouse retina, for example, become more linear with decreasing light intensity (Grimes et al., 2014).

Since we presented natural images in a full-field fashion, we furthermore need to consider RF surround effects that likely also contribute to differences between cells in natural stimulus encoding. A study in the primate retina, for example, showed that surround activation can alter spatial integration in the RF center (Turner et al., 2018). We cannot exclude that disregarded surround effects contributed to shortcomings of the LN model, but we tried to minimize confounding consequences of surround activation. First, our LN model could contain inhibitory surround contributions, because the model’s linear filter was a difference of Gaussians fitted to the spike-triggered average, which could contain positive and negative values. This might not reflect the actual surround strength, however, which could be underestimated with spatiotemporal white noise (Wienbar and Schwartz, 2018). Yet, it should reduce the risk that neglected surround effects obscure spatial-integration effects on LN model performance. In line with this, our spatial contrast sensitivity analysis showed that we could partially explain across-cell differences in LN model performance by only considering local contrast in the RF center. And likewise, rectification indices obtained from the full-field and from the local checkerboard flash stimulus worked equally well to explain the performance differences of the LN model, corroborating that nonlinear integration in the center is a main factor in LN model performance. Finally, one should note here that RF surround strength may be a strong covariate of nonlinear spatial integration in the mouse retina, as many strongly nonlinear cell types also show strong surround suppression (Jacoby and Schwartz, 2017; Mani and Schwartz, 2017; Zhang et al., 2012).

### Implications for neuronal modeling

Proposed improvements to LN-type models go in many directions (Latimer et al., 2019; Shi et al., 2019). Here we demonstrated that the incorporation of sensitivity to fine spatial structure into models (e.g. with spatial subunits) should be significant for natural stimuli. We found that cells with low LN model performance mostly showed nonlinear spatial integration and that rectification of non-preferred contrasts in the RF center was particularly important. This observation of the importance of rectification agrees with results from nonlinear subunit modelling of ganglion cells: in the macaque retina the rectification of subunits determines the degree of nonlinear integration under natural images (Turner and Rieke, 2016), and in a model of salamander retinal ganglion cells under white-noise stimulation, threshold-linear rectification of subunit signals worked nearly as well as more elaborate, fitted shapes (Real et al., 2017). The checkerboard flash stimuli we developed can be used to efficiently estimate the degree of subunit rectification for many retinal ganglion cells simultaneously. Given the recently developed techniques for estimating subunit locations (Liu et al., 2017; Maheswaranathan et al., 2018; Shah et al., 2020), this paves the way for building more detailed models for different ganglion cell types. Our results also indicate that such cell-type specific approaches may be needed as there might not be a satisfactory single “standard model” (Carandini et al., 2005).

## Acknowledgements

We thank Michael Weick for help in designing experiments and for discussion. This work was supported by the Boehringer Ingelheim Fonds, the European Research Council (ERC) under the European Union’s Horizon 2020 research and innovation programme (grant agreement number 724822), and by the Deutsche Forschungsgemeinschaft (DFG, German Research Foundation) – Projektnummer 154113120 - SFB 889, project C1).

## Author Contributions

Conceptualization: D.K and T.G., Methodology: D.K. and T.G., Software: D.K., Formal Analysis: D.K., Writing – Original Draft: D.K., Writing – Review & Editing: D.K. and T.G., Visualization: D.K., Supervision: T.G., Funding Acquisition: T.G., Resources: T.G.

## Competing interests

The authors declare no competing interests.

## Materials and Methods

### Animals

We used 13 retina pieces from 9 adult wild-type mice of either sex (6 C57BL/6J and 3 C57BL/6N, 7 male and 2 female), mostly between 8-12 weeks old (except for one 18- and one 26-week-old). All mice were housed in a 12-hour light/dark cycle. Experimental procedures were in accordance with national and institutional guidelines and approved by the institutional animal care committee of the University Medical Center Göttingen, Germany.

### Tissue preparation and electrophysiology

Mice were dark-adapted for at least an hour before eye enucleation. After the animal had been sacrificed, both eyes were removed and immersed in oxygenated (95% O_2_-5% CO_2_) Ames’ medium (Sigma-Aldrich), supplemented with 22 mM NaHCO_3_ (Merck Millipore) and 6 mM D-glucose (Carl Roth). We cut the globes along the ora serrata, removing the cornea, lens, and vitreous humor. The resulting eyecups were often cut in half to allow two separate recordings. Before the start of each recording, we isolated retina pieces from the eyecups. We placed the pieces ganglion cell-side-down on planar multielectrode arrays (Multichannel Systems; 252 electrodes; 30 μm diameter, either 100 or 200 μm minimal electrode distance) with the help of a semipermeable membrane, stretched across a circular plastic holder (removed before the recording). The arrays were coated with poly-D-lysine (Merck Millipore). Throughout the recording, retinal pieces were continuously superfused with the oxygenated Ames solution flowing at ∼250 ml/h. The bath solution was heated to a constant temperature of 34–35°C via an inline heater in the perfusion line and a heating element below the array. Dissection and mounting were performed under infrared light on a stereo-microscope equipped with night-vision goggles.

Extracellular voltage signals were amplified, band-pass filtered between 300 Hz and 5 kHz, and digitized at 10 kHz sampling rate. Spikes were detected by threshold crossings (four standard deviations of the voltage trace), and spike waveforms were sorted offline into units with a custom-made IgorPro (WaveMetrics) routine based on Gaussian mixture models (Pouzat et al., 2002). We curated the routine’s output and selected only well-separated units with clear refractory periods. Duplicate units were identified by temporal cross-correlations and removed. Finally, only units with stable electrical images (Litke et al., 2004) throughout the recording were considered for further analysis.

### Visual stimulation

Visual stimuli were generated and controlled through custom-made software, based on Visual C++ and OpenGL. Different stimuli were presented sequentially to the retina through a gamma-corrected monochromatic white OLED monitor (eMagin) with 800×600 square pixels and 60 Hz refresh rate.

The monitor image was projected through a telecentric lens (Edmund Optics) onto the photoreceptor layer of the retina, and each pixel’s side measured 7.5 μm on the retina. All stimuli were presented on a background of low photopic light levels (2.5 or 3.5 mW/m^2^, corresponding to 6.3×10^3^ or 7.9×10^3^ R*/rod/s), and their mean intensity was always equal to the background. We fine-tuned the focus of stimuli on the photoreceptor layer before the start of each experiment by visual monitoring through a light microscope and by inspection of spiking responses to contrast-reversing gratings with a bar width of 30 μm.

### Linear receptive field identification

To estimate the RF of each cell, we used a spatiotemporal binary white-noise stimulus (100% contrast) consisting of a checkerboard layout with flickering squares (60 μm side). The update rate was either 30 or 60 Hz in different experiments. We measured the spatiotemporal RF by calculating the spike-triggered average (STA) over a 500-ms time window (Chichilnisky, 2001), and fitted a parametric model to the RF (Chichilnisky and Kalmar, 2002). The model was spatiotemporally separable and comprised a product of a spatial (*k*_*S*_(***x***)) and a temporal component (*k*_*T*_(*t*)).

The spatial component was modelled as a difference of Gaussians:

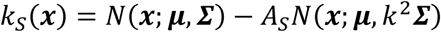

where 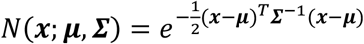 is a two-dimensional Gaussian function with mean ***μ*** and covariance matrix ***Σ*** (describing the RF center’s coordinates and shape), *A*_*s*_ ∈ [0,1] captures the RF surround strength relative to the RF center, and *k* ≥ 1 is a scaling factor for the surround’s extent.

The temporal component was modelled as a difference of two low-pass filters:

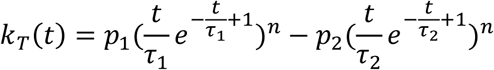

with *t* > 0 indicating the time before the spike and *p*_1_ > 0, *p*_2_ > 0, *τ*_1_ > 0, *τ*_2_ > 0, *n* > 0 being free parameters.

We fitted the full parametric model (*k*_*S*_(***x***) · *k*_*T*_(*t*)) to the STA by minimizing the mean squared error using constrained nonlinear optimization. To get reasonable initial conditions, we first separately fitted the spatial component to the STA frame at which the element with the largest absolute value occurred and the temporal component to the time course of the same element. If the element was negative, the sign of the STA frame was inverted before the spatial component fit. We then seeded the obtained values of spatial and temporal fits as the initial parameters for the full spatiotemporal fit. The chosen initialization procedure caused *k*_*S*_ to always have a positive center peak, thus resolving the ambiguity in the signs of *k*_*S*_ and *k*_*T*_.

Cells were classified as either ON- or OFF-type based on the sign of the first peak (i.e. closest to zero) in the fitted temporal component *k*_*T*_(*t*). Here we disregarded a peak if its amplitude (unsigned) was smaller than 25% of the largest deflection. The diameter of the RF center was defined as the diameter of a circle with the same area as the 2σ (elliptical) boundary of the Gaussian center profile (Baden et al., 2016). We also used the 2σ boundary for all RF center visualizations.

### Natural image response predictions with a linear-nonlinear model

We selected natural images as stimuli from three sources: the van Hateren Natural Image Dataset (van Hateren and van der Schaaf, 1998), the McGill Calibrated Colour Image Database (Olmos and Kingdom, 2004) and the Berkeley Segmentation Dataset (Arbeláez et al., 2011). The central square region of each image was resized to 512×512 pixels (400×400 pixels in a few experiments) by cropping (van Hateren and McGill images) or cropping and upsampling with nearest neighbor interpolation (Berkeley images). All color images (McGill and Berkeley databases) were converted to grayscale by weighted averaging over the color channels (Liu et al., 2017). We normalized the mean and standard deviation of the pixel values for each image by appropriately shifting and scaling the values so that the mean pixel intensity was equal the background and the standard deviation was 40% of the mean intensity. Pixel values that then deviated from the mean by more than 100% in either direction were clipped to ensure that the maximal pixel values were within the physically available range of the display. Finally, all images were encoded at 8-bit color depth, to match the range of our OLED monitor. The images were presented on top of a uniform grey background and centered on the multielectrode array, covering a region of 3.84×3.84 mm^2^ on the retina (3×3 mm^2^ for 400×400 pixels).

In every experiment, we used 300 natural images (100 from each database), except for one (200 images in total). Images were presented individually for 200 ms each, with an 800-ms inter-stimulus-interval of homogeneous background illumination. We collected ten trials for each image, by consecutively presenting ten different pseudo-randomly permuted sequences of all images. For each cell, we measured the response as the trial-averaged number of spikes over a 250-ms window following stimulus onset.

To compare a cell’s responses to model predictions, we constructed a linear-nonlinear (LN) model (Chichilnisky, 2001), which generates average spike count responses *R*_*m*_ ≥ 0 to natural image stimuli ***s***_*m*_: *R*_*m*_ = *f*(***k***^***T***^ · ***s***_*m*_), where the vector ***k*** is a linear spatial filter, *f* is a nonlinear function and *m* denotes the image index. For the analyses, all natural image stimuli were spatially clipped to the smallest square that could fit the 4σ boundary of the RF center, and their pixel intensity values were transformed to Weber contrast values, which constitute the elements of ***s***_*m*_. For the linear filter (***k***), we used the parametric spatial RF component (*k*_*S*_) estimated from white noise, sampled at the center point of each pixel of the clipped natural image. The linear filter was normalized to unit sum of the absolute values of its elements. Linear predictions (*g*) were estimated from the inner product of stimuli and the linear filter: *g*_*m*_ = ***k***^***T***^ · ***s***_*m*_. Because the linear filter is composed of mainly positive values, the sign of the linear prediction reflects the net contrast in the spatial RF. For the nonlinear part of the LN model (*f*) we used a bi-logistic nonlinearity of the form

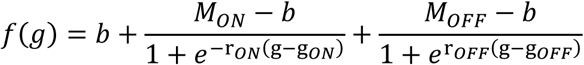

where *b* ≥ 0, *M*_*ON*_ ≥ *m, M*_*OFF*_ ≥ *m, r*_*ON*_, ≥ 0, *r*_*OFF*_ ≥ 0, *g*_*ON*_, *g*_*OFF*_ were free parameters that were fitted to data. To facilitate estimation of the parameters for both monotonic and U-shaped nonlinearities, we first fitted a single logistic nonlinearity 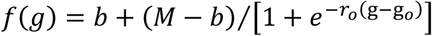 to the data, and initialized the parameters of the bi-logistic nonlinearity to describe the dominant lobe (ON or OFF, determined by the sign of *r*_*o*_). Such bi-logistic nonlinearities had been previously used to describe tuning curves in sensory neuroscience, under the name “difference of sigmoids” (Fischer et al., 2009; Mel et al., 2018; Murgas et al., 2019).

To assess the prediction accuracy by the LN model, we applied a normalized correlation coefficient (CC_norm_) as our model performance metric (Schoppe et al., 2016). This measure was used in order to account for differences in response reliability among cells, since we used a relatively small number of trials per image. Concretely, for *M* images and *N* trials per image, *R*_*m,n*_ denoted the cell’s response to image *m* for trial *n*, with 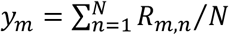 being the average response to a particular image, 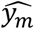 being the prediction for the same image, and with *y* and *ŷ*denoting the corresponding distributions over images. CC_norm_ was then defined as

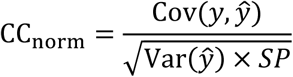

We calculated the necessary quantities as the sample covariance 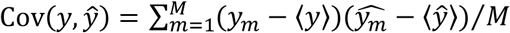 and variance 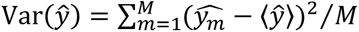. *SP* denotes the signal power, a measure of signal-to-noise ratio, and is defined as

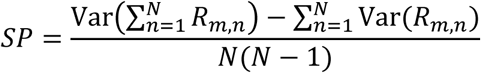

with the two variances in the numerator denoting sample variances over images.

We estimated LN model performance through ten-fold cross-validation. Briefly, the collection of average responses for all images was randomly split into ten equally-sized sets. Every set was used once as a test set for the full LN model, whose nonlinearity was fitted to the other 90% of image responses. For each cell, LN model performance was defined as the average CC_norm_ over all cross-validation sets. For all nonlinearity visualizations in the plots, we used the nonlinearity corresponding to the cross-validation set with the CC_norm_ value closest to the average.

Since CC_norm_ values are ill-defined for very low data reliability, we excluded cells whose responses for identical images were highly variable. For each cell, we therefore calculated the coefficient of determination (R^2^) between responses averaged over even 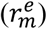 and over odd trials 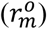, where m = 1,…,M enumerates the images. Concretely, we used a symmetrized R^2^, defined as

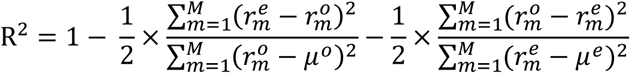

where *μ*^*o*^ and *μ*^*e*^ are the average odd and even trial responses over images. We excluded cells with R^2^<0.5 from further analysis. This criterion included 959 out of 1209 recorded cells in the analysis.

### Calculation of spatial contrast sensitivity for natural images

We measured the spatial contrast (SC) of an image in the RF center of a given ganglion cell as the weighted standard deviation of pixel contrast values inside the 2σ contour of the Gaussian center fit:

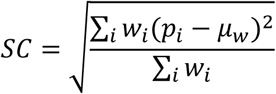

where the sums run over all pixels *i* within the 2σ contour, *p*_*i*_ is the pixel value, *w*_*i*_ is the pixel weight as given by the value at the pixel center of the fitted RF center part, and *μ*_*w*_ is the weighted mean of the pixel values.

To obtain the spatial contrast sensitivity, we sorted the images according to their linear predictions in the cell’s LN model and then grouped neighboring images into pairs, with each image belonging to a single pair, yielding 150 pairs per cell when 300 images had been applied. For each image pair, we calculated the spatial contrast difference and the trial-averaged response difference. To compare across cells, we normalized the response differences by the maximum response (over images) of the cell. We defined the spatial contrast sensitivity as the slope of the linear regression between the spatial contrast differences and the normalized response differences. Cells were defined as contrast sensitive if they had a significant regression slope at the 5% significance level.

### Assessment of spatial nonlinearity with contrast-reversing gratings

To compare our findings to classical analyses of spatial integration, we stimulated the retina with full-field square-wave gratings of 100% contrast. The contrast of the gratings was reversed every 1 s. The reversing gratings were presented sequentially from higher to lower spatial frequencies for 20-30 reversals each, and the whole sequence was repeated two times. Depending on the experiment, we sampled five to eight spatial frequencies, with bar width ranging from 15 to 240 μm. For each spatial frequency, we applied one to four equidistant spatial phases, with more phases for lower spatial frequencies (e.g. one for 15, two for 30, two for 60, four for 120, four for 240 μm bar width). In some of the recordings, we also included contrast reversals of homogeneous illumination (corresponding to a bar width of 6 mm or larger). Between presentations of the different gratings, there was a grey screen at background intensity for 2 s. We constructed PSTHs over one reversal period by binning ganglion cell spikes with 10-ms bins and averaging across reversals and repeats, leaving out the first reversal after a gray period. In order to exclude cells with unreliable responses, we calculated R^2^ values between average response vectors of even and odd trials, similar to the analysis of natural-image responses. We created the response vector of a single trial by concatenating single-trial PSTHs from all different spatial frequencies and phases. We only considered cells with R^2^>0.1 for our population analyses. The criterion was satisfied by 890 out of 1126 cells recorded for this stimulus.

To estimate the *grating* spatial scale for each cell, we extracted the peak firing rate in the PSTH (across time and spatial phases) for each bar width (Krieger et al., 2017). We then fitted a logistic function (cf. Natural image response predictions) to the relationship of peak rates versus bar width, and extracted the function’s midpoint as an estimate of the spatial scale. The amplitudes of harmonics of the PSTH were calculated by temporal Fourier transforms for each combination of spatial frequency and phase (Hochstein and Shapley, 1976). From the PSTHs of all spatial scales and phases, we extracted the maximum amplitude F1 at the stimulus frequency as well as the maximum amplitude F2 at twice the stimulus frequency and defined the nonlinearity index as the ratio of F2 over F1. This definition is slightly different than other approaches, where the F2/F1 ratio is calculated for each spatial scale and phase separately, with the maximum being chosen as the nonlinearity index (Carcieri et al., 2003; Hochstein and Shapley, 1976; Petrusca et al., 2007). Our approach aimed at capturing the maximum mean-luminance-induced modulation in F1 and the maximum spatial-contrast-induced modulation in F2.

### Assessment of spatial input nonlinearities with checkerboard flashes

To assess how local visual signals are transformed in nonlinear cells, we used a stimulus that had a checkerboard layout with square tiles of either 105 or 120 µm to the side. The tiles were alternatingly assigned to two sets (A and B) so that neighboring tiles were in different sets. For each individual stimulus presentation, each set of tiles was assigned an intensity *s*_*A*_ or *s*_*B*_, respectively, expressed as the Weber contrast from background illumination. Similar to our presentation of natural images, these checkerboard stimuli were flashed for 200 ms with an inter-stimulus interval of 800 ms, during which background illumination was presented. The contrast pairs (*s*_*A*_, *s*_*B*_) were selected from a two-dimensional stimulus space organized in polar coordinates, by using 24 equidistant angles, each with 10 equidistant radial contrast values 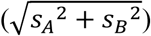 between 3-100% (see Figure 4A, bottom-right), and presented in pseudorandom order. The set of all contrast pairs was presented to the retina 4 to 5 times, with a different pseudorandomly permuted sequence chosen each time. We calculated cell responses by counting the number of spikes for each ganglion cell over a 250-ms window following stimulus onset and averaging over trials. Iso-response contour lines were constructed from the cells’ response profiles using MATLAB’s *contour* function. To exclude cells with unreliable responses, we calculated R^2^ values across the set of all contrast combinations between spike counts averaged over even and over odd trial numbers. We only considered cells with R^2^>0.1 for our population analyses. This criterion was satisfied by 833 out of 1204 cells for which the stimulus was recorded.

Rectification (RI) and convexity (CI) indices were calculated for a specific contrast level, here *c* = 0.***6***. To quantify rectification of non-preferred contrasts (RI), we compared the responses *r*^*half*^ under stimulation with only one spatial input (e.g. *s*_*A*_ = *c* and *s*_*B*_ = 0, corresponding to a stimulus on one of the four half axes of the stimulus space) to the responses *r*^*oppos*^ under stimulation with this input and the other spatial input at opposite contrast (*s*_*A*_ = *c* and *s*_*B*_ = −*c*). In the cases with no direct response measurement for a particular contrast pair, we estimated the response based on the measured responses to nearby contrast pairs, using natural neighbor interpolation, as implemented in MATLAB’s *scatteredInterpolant* function. From all response measurements, we subtracted the background spike count, measured as the response to the (0, 0) pair, which was included as a regular stimulus in the sequence of contrast pairs. To use a single definition of RI for ON, OFF, and ON-OFF cells, we considered all four half axes in the stimulus space (with either *s*_*A*_ or *s*_*B*_ at either positive or negative contrast) and computed a weighted average from the four *r*^*half*^ values as well as from the corresponding *r*^*oppos*^ values (note that there are only two *r*^*oppos*^ values that are each used twice) to define RI as their ratio:

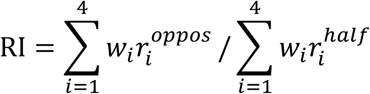

where the weights *w*_*i*_ are measures of sensitivity along each half axis *i*. Concretely, we obtained *w*_*i*_ as the slope of a regression line, fitted to the contrast-response pairs along the corresponding half axis.

Similarly, for quantifying integration of preferred contrasts (CI), we compared the *r*^*half*^ values to responses *r*^*same*^, which were measured with the same contrast for the two stimulus components, *s*_*A*_ = *s*_*B*_ = *c****/***2, corresponding to the spatially homogeneous stimulus that has the same linearly integrated contrast as the stimulus used to measure *r*^*half*^. Again, we took all four half-axes into account for defining CI:

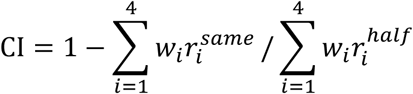

We subtracted the ratio of *r*^*same*^ over *r*^*half*^ from unity so that CI = 0 corresponds to linear integration and CI > 0 to *r*^*same*^ < *r*^*half*^, and thus a convex, outward-bulging shape of the iso-response contour line. We used rectification and convexity indices to formally define homogeneity-sensitive cells as cells with RI < 0 and CI < 0, corresponding to iso-response contour lines curving towards the origin.

To probe spatial integration in the RF center with minimal surround influence, we used local checkerboard flashes. The stimulus was similar to the one above, but with small patches of 2×2 tiles. Tiles here had a side length of 105 μm, and patches thus had a side length of 210 μm. To compare our results with the full-field version of the stimulus, patch tiles were placed to align with the tiles of the full-field stimulus. The local patches were flashed for 200 ms, with no interval between successive presentations. For each individual presentation, the applied patch locations were randomly chosen to maximally fill the screen (typical number of locations = 44-61, median = 53), while ensuring a minimum center-to-center distance of three patch side lengths (630 µm) for simultaneously presented patches (Figure 4C, bottom-left). The rest of the screen was kept at the background illumination (Figure 4C, top). For each presented patch, the contrast combination (*s*_*A*_, *s*_*B*_) was selected randomly and independently from the contrast combinations at other, simultaneously displayed locations. We applied fewer contrast combinations than for the full-field version of the stimulus to ensure adequate numbers of trials for each contrast combination at each location. Specifically, we used 8 or 12 equidistant angles in the stimulus space, each with 5 or 6 equidistant radial values between either 20-100% or 3-100%.

For analysis, we selected for each ganglion cell the patch closest to its RF center and extracted the responses to flashes when this particular patch was used. We counted the number of spikes over a 250-ms window following presentation onset and again subtracted the background activity, which was here obtained by interpolation to the (0, 0) contrast pair. Response contour lines in stimulus space as well as rectification and homogeneity indices were calculated in the same way as for the full-field version of the checkerboard flashes. Similarly to the full-field stimulus, we calculated R^2^ values between the average spike counts of even and odd trials with respect to all contrast combinations, and only considered cells with R^2^>0.1 for our population analyses. This criterion was satisfied by 399 out of 562 cells.

### Spatial scale estimation from blurred natural images

For recordings with blurred natural images, we selected either 30 or 40 images from our set of natural images. The images were blurred by convolution with a two-dimensional, spherically symmetric Gaussian function. We used different σ-values of the Gaussian to implement different spatial scales of blurring, defined as the diameter of the 2σ Gaussian contour (Schwartz et al., 2012), to also match our RF center definition. Blurred and original images were presented in a pseudorandom sequence, similar to the presentation of the large set of natural images described above, collecting 10 trials for each image and blurring scale. Responses were again measured for each ganglion cell by counting the number of spikes over a 250-ms window following stimulus onset.

We calculated R^2^ values (see Natural image response predictions) between blurred and original spike counts for each scale. We also calculated an R^2^ value for the original image responses, by considering odd- and even-trial averages and assigned this value to a blurring scale of 0 μm. We then fitted logistic functions to the R^2^ values with respect to the blurring scales. We defined the natural spatial scale for each ganglion cell as the midpoint of the fitted logistic function. Again, by requiring R^2^>0.1 for odd-versus even-trial averages of the original images, we included 747 cells out of 850 for which we had recorded the stimulus.

### Detection of image-recurrence-sensitive cells

We detected image-recurrence-sensitive (IRS) cells as described previously (Krishnamoorthy et al., 2017). Briefly, we presented a square-wave grating of either 240 or 270 µm spatial period and 60% contrast in a sequence of 800-ms-long fixations, separated by 100-ms transitions. During a transition, the grating was shifted by approximately two spatial periods to land in one of four equidistant fixation positions (corresponding to four specific spatial phases of the grating). The sequence of the four fixation positions was randomly chosen so that all 16 possible transitions (between starting and target positions) appeared several times in the stimulus sequence.

IRS cells are described as cells that show a strong response peak after onset of the new fixation when the grating position is the same as before the transition, but not when it has reversed contrast across the transition. To detect this, as done previously (Khani and Gollisch, 2017; Krishnamoorthy et al., 2017), we measured the response after each of the 16 possible transitions by creating PSTHs with bins of 10 ms and extracting the maximal difference for successive time bins in the PSTH as a measure of response increase (maximal derivative of the PSTH) in the window from 50 to 200 ms after fixation onset. We compared for each target grating *i* the maximal derivative 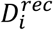 under image recurrence (when the starting grating was also *i*) to the maximal derivative 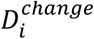 when the starting grating was contrast-reversed compared to grating *i*. We calculated a recurrence sensitivity index (*RSI*) as 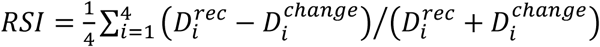. Cells with RSI>0.7 and an average peak firing rate of at least 50 Hz in the post-transition PSTHs of the four image recurrences were considered as IRS cells.

### Detection of direction- and orientation-selective cells

To identify direction-selective ganglion cells, we used drifting sinusoidal gratings of 100% contrast, 240 µm spatial period, and a temporal frequency of 0.6 Hz (Sabbah et al., 2017). The gratings were shown in a sequence of eight equidistant directions with four temporal periods per direction, separated by 5 s of background illumination. The sequence was repeated four to five times. For each angle (*θ*), we collected the average spike responses (*r*_*θ*_) during the presentation of the grating (excluding the first period). We calculated a direction selectivity index (DSI) as the magnitude of the normalized complex sum ∑_*θ*_ *r*_*θ*_*e*^*iθ*^***/***∑_*θ*_ *r*_*θ*_ (Mazurek et al., 2014). The preferred direction was obtained as the argument of the same sum.

We also used drifting square-wave gratings of 100% contrast, 225 µm spatial period, and a temporal frequency of 4 Hz to identify orientation-selective ganglion cells (Nath and Schwartz, 2016, 2017). The gratings were shown in a sequence of eight equidistant directions with twelve periods per direction, separated by 2 s of background illumination. The sequence was repeated four to five times. We calculated an orientation selectivity index (OSI) as the magnitude of the complex sum ∑_*θ*_ *r*_*θ*_*e*^*i*2*θ*^***/***∑_*θ*_ *r*_*θ*_. The preferred orientation was obtained as the line perpendicular to half the argument of the same sum.

To calculate the statistical significance for both indices, we used a Monte Carlo permutation approach (Liu et al., 2017). For a given cell, we repeatedly shuffled the responses over all angles and trials 2000 times to obtain a distribution of DSI (or OSI) values under the null hypothesis that the firing rates are independent of the motion direction (or orientation). All cells with DSI>0.25 (significant at 1% level) where considered as DS cells. Similarly, OS cells were identified as cells with OSI>0.25 (significant at 1% level) that were not DS or IRS. We only included cells with a total mean firing rate >1 Hz during the presentation of the drifting gratings (Kühn and Gollisch, 2016).

### Statistical testing

For all statistical procedures, we used the default MATLAB2018b functions. For calculating correlations between various cell-specific measures, such as LN model performance or spatial contrast sensitivity, we used a non-parametric correlation coefficient (Spearman’s ρ) instead of the linear Pearson’s r. For comparing dependent and independent sample measurements, we used Wilcoxon sign-rank and rank-sum tests, respectively.

### Data and Code Availability

Example cell data and analysis code will be made available upon publication.

## Supplemental Figures

**Figure S1.**
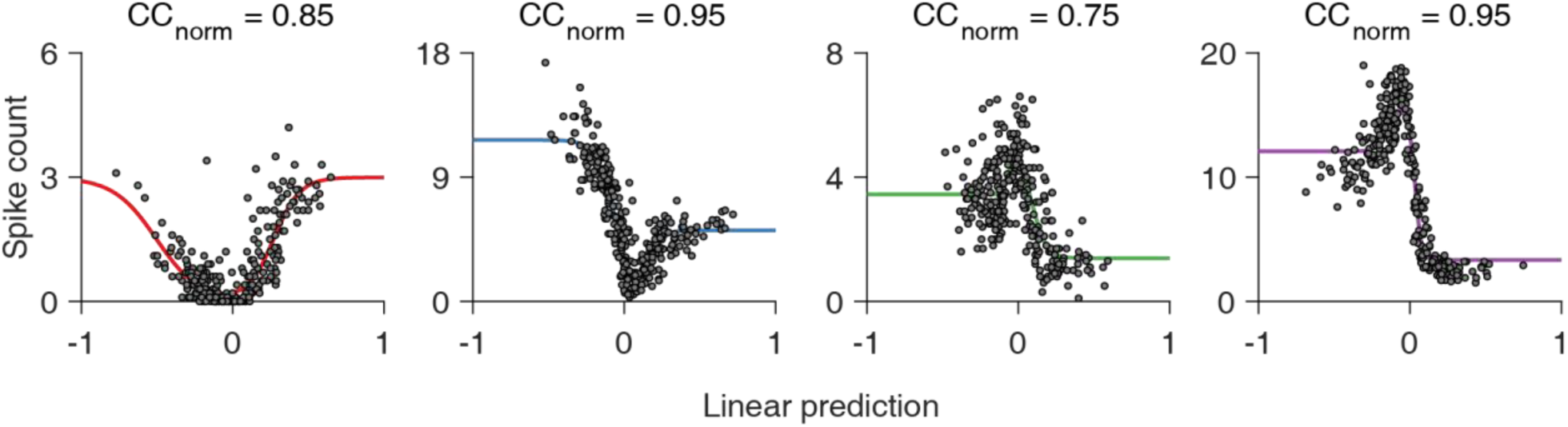
Examples of U- and bell-shaped nonlinearities. Four sample cells, with either ON-OFF-type nonlinearities (the two leftmost), or contrast-suppressed-type nonlinearities (the two on the right).

**Figure S2.**
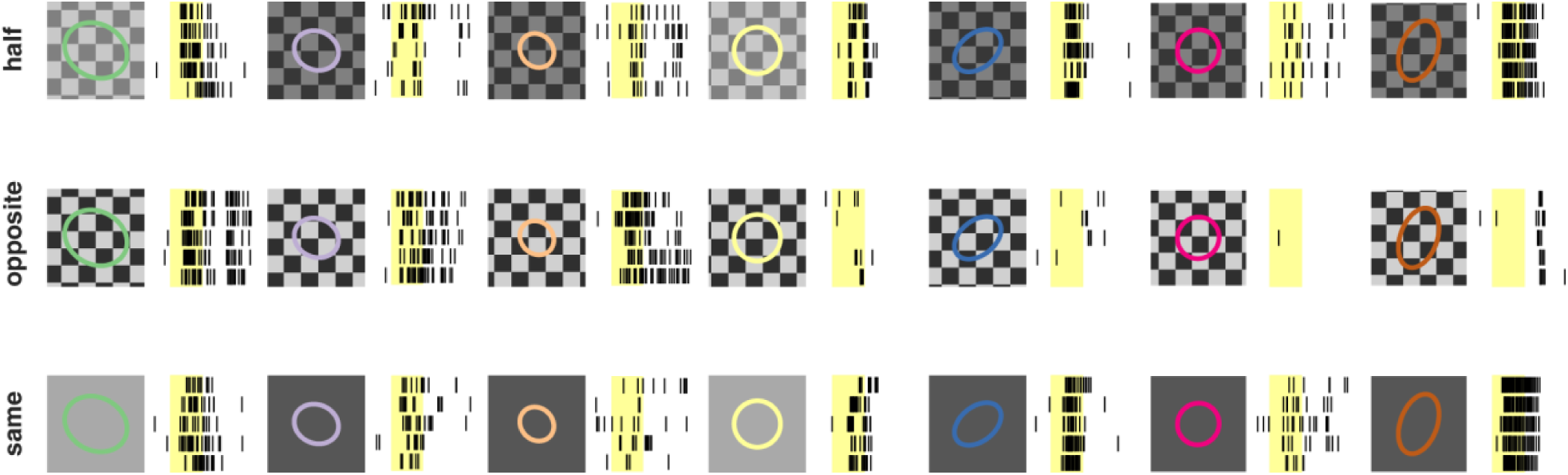
Responses to contrast pairs used for measuring rectification and convexity indices. Each column with stimulus and receptive field (RF) on the left and corresponding raster plots (five trials) on the right corresponds to one cell. Yellow shaded regions mark the 200-ms image presentations. All seven cells match the corresponding ones from Figure 4, following the same order and having their RF outlines colored similarly.

**Figure S3.**
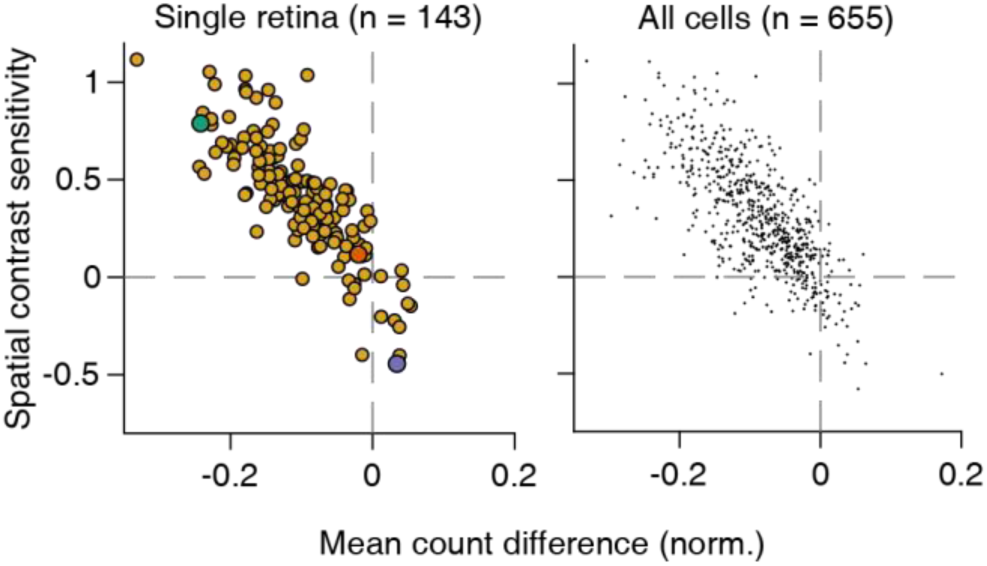
Relationship between spatial contrast sensitivity and spike count difference induced by blurring of natural images. Left: single retina (Spearman’s ρ = −0.83, p < 10^−37^), differently colored points match the corresponding cells from Figure 6. Right: pooled ganglion cell population (Spearman’s ρ = −0.73, p < 10^−112^) from 9 retinas, 6 animals. Dashed lines in both panels show values indicating linear spatial integration.

**Figure S4.**
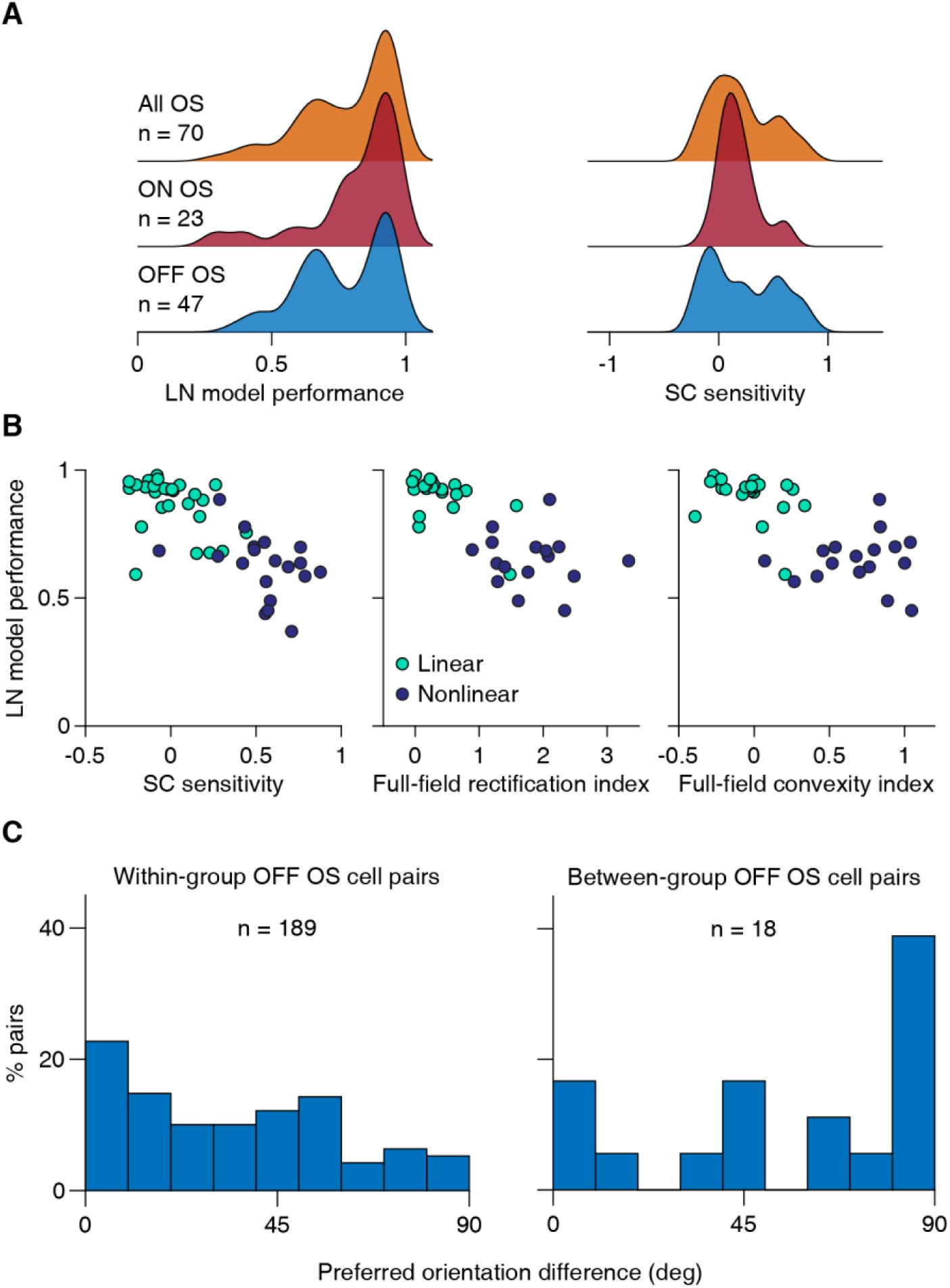
Subtypes of orientation-selective (OS) ganglion cells differ in their spatial integration features. **A)** Distributions of LN model performance and spatial contrast (SC) sensitivity for ON and OFF OS cells in comparison to all OS ganglion cells. ON cells show rather linear spatial integration, whereas OFF cells had a bimodal distribution. **B)** OFF OS cells were assigned to two groups (linear and nonlinear) with k-means clustering. The features we used were the LN model performance, spatial contrast sensitivity, full-field rectification and convexity indices. All four features were available for only n = 39/47 OFF OS cells, and clustering was performed with these cells only. For the other cells, responses to checkerboard flashes were not recorded, and they were assigned to the group whose cluster centroid was closest for the two available measures, LN model performance and spatial contrast sensitivity. **C)** To examine whether the preferred orientations of linear vs nonlinear OFF OS cells systematically differed, we compared the distributions of differences in preferred orientation for pairs of OS cells that belonged to either the same group (“within-group”) or to different groups (“between-group”). The differences in preferred orientation were significantly smaller for within-group pairs than for between-group pairs (Wilcoxon rank-sum test, p = 0.0056).

## Notes

### Competing Interest Statement

The authors have declared no competing interest.

